# Altered DNA methylation underlies monocyte dysregulation and innate exhaustion memory in sepsis

**DOI:** 10.1101/2023.08.30.555580

**Authors:** Blake A. Caldwell, Yajun Wu, Jing Wang, Liwu Li

## Abstract

Innate immune memory is the process by which pathogen exposure elicits cell-intrinsic states to alter the strength of future immune challenges. Such altered memory states drive monocyte dysregulation during sepsis, promoting pathogenic behavior characterized by pro-inflammatory, immunosuppressive gene expression in concert with emergency hematopoiesis. Epigenetic changes, notably in the form of histone modifications, have been shown to underlie innate immune memory, but the contribution of DNA methylation to this process remains poorly understood. Using an *ex vivo* sepsis model, we discovered broad changes in DNA methylation throughout the genome of exhausted monocytes, including at several genes previously implicated as major drivers of immune dysregulation during sepsis and Covid-19 infection (e.g. *Plac8*). Methylome alterations are driven in part by Wnt signaling inhibition in exhausted monocytes, and can be reversed through treatment with DNA methyltransferase inhibitors, Wnt agonists, or immune training molecules. Importantly, these changes are recapitulated in septic mice following cecal slurry injection, resulting in stable changes at critical immune genes that support the involvement of DNA methylation in acute and long-term monocyte dysregulation during sepsis.

## INTRODUCTION

In the innate immune system, monocytes carry the essential role of immune sentinels, migrating through the bloodstream to sites of tissue damage or infection and orchestrating the initial immune response through cytokine release and phagocytosis of infectious cells and molecules (Turvey & Broide, 2010; Geissmann et al, 2010; Sica & Mantovani, 2012). Dynamic control of this process is achieved through innate immune memory (IM), in which monocyte exposure to injury or pathogen-associated molecular patterns (PAMPs) results in the functional reprogramming of these cells to alter their response to future stimulation (Quintin *et al*, 2014). IM pathways can be specific to a given PAMP through the activity of different pattern recognition receptors (PRRs) and are broadly classified into two functionally opposed mechanisms: training and tolerance (Netea *et al*, 2020a). In immune training, PAMP exposure elicits a heightened immune state of improved PRR pathogen recognition and enhanced inflammatory activity in response to subsequent immune challenges (Netea *et al*, 2011). By contrast, in immune tolerance, PAMP exposure dampens a cell’s inflammatory response to future immune challenges through the silencing of pro-inflammatory genes. This IM mode has been exemplified by the lipopolysaccharide (LPS) tolerance model, in which repeated exposure to the bacterial endotoxin LPS inhibits the expression of inflammatory cytokines through NF-κB signaling (Yan et al, 2012; Foster et al, 2007; Dobrovolskaia & Vogel, 2002; Fan & Cook, 2016).

Monocyte exhaustion represents a pathogenic form of IM in which prolonged PAMP exposure elicits a paradoxical state defined by both pro-inflammatory and immunosuppressive gene expression in concert with emergency hematopoiesis through cellular de-differentiation. (Pradhan *et al*, 2021). This memory state, suggestive of a failure of emergency feedback mechanisms, is characteristic of the “cytokine storm” phenomenon commonly observed in sepsis patients and severe Covid-19 infection (Xiao *et al*, 2011; Tang *et al*, 2020; Hu *et al*, 2021). However, despite the tremendous burden of these disorders on global healthcare at the levels of both patient mortality and treatment cost, the molecular underpinnings of monocyte exhaustion, as well as the sustained immunoparalysis of its survivors, remain poorly understood. (Baudesson de Chanville et al, 2020; Gentile et al, 2012; Prescott & Angus, 2018).

Increasing evidence suggests that epigenetics, or changes to chromatin above the level of DNA sequence, function as the major driver of transcriptional reprogramming in IM (Smale *et al*, 2014; Rodriguez *et al*, 2019; Netea *et al*, 2020a). In a landmark study conducted through the BLUEPRINT Consortium, it was demonstrated that genome-wide changes in active histone marks H3K27ac and H3K4me1/3 underlie many of the gene expression changes observed during monocyte-to-macrophage differentiation under immune tolerance and training conditions, corroborating the results of earlier studies focusing on individual genes (Saeed *et al*, 2014; Foster *et al*, 2007; Kleinnijenhuis *et al*, 2012; Bekkering *et al*, 2014; De Santa *et al*, 2007, 2009). As a result, much subsequent research into the epigenetic regulation of IM has focused on the contribution of histone modifications to this process. However, epigenetic regulation is a complex interplay of numerous chromatin features, including nucleosomal remodeling, long non-coding RNA interactions, and DNA methylation in the form of 5- methylcytosine (5mC) (Zhang *et al*, 2020). Despite evidence that epigenetic regulation through DNA methylation may play a role in immune tolerance during macrophage differentiation and inflammatory signaling of peripheral blood monocytes in septic patients, the contribution of 5mC to IM is critically underexplored (Novakovic et al, 2016; Lorente-Sorolla et al, 2019).

In this study, we utilized our lab’s recently published *ex vivo* mouse sepsis model to identify DNA methylation changes that occur during monocyte exhaustion and evaluate their regulatory significance as drivers of the exhaustion phenotype. In this manner, we identified thousands of differentially methylated regions in exhausted monocytes and linked their alteration to transcriptional changes at critical immune genes. Utilizing predictive modeling based on a monocyte single-cell RNA sequencing (scRNA-seq) dataset, we were next able to implicate the Toll-like receptor 4 (TLR4) signaling component TICAM2 and Wnt signaling as upstream regulators of DNA methylation changes observed in exhausted monocytes. Furthermore, we demonstrate the therapeutic benefits of DNA methyltransferase (DNMT) inhibitor 5-azacytidine (5-aza) or immune adjuvant trehalose 6,6’-dimycolate (TDM) in limiting the extent of DNA methylation changes in exhausted monocytes and restoring immune function. Finally, we were able to validate DNA methylome reprogramming in bone marrow monocytes of septic mice following cecal slurry injections, demonstrating long-term 5mC changes at key immune genes. Taken together, these findings highlight the underappreciated role DNA methylation plays in regulating pathogenic monocyte behavior, as well as implicate DNA methylation as a potential target for therapeutic intervention in disorders characterized by monocyte dysregulation.

## RESULTS

### DNA methylation reprogramming at gene regulatory elements in exhausted monocytes

DNA methylation in the form of 5-methylcytosine (5mC) at gene promoters and enhancers functions as a critical regulator of cell state-specific gene expression profiles, most commonly through transcriptional silencing (Bird, 2007; Smith & Meissner, 2013). In order to test the involvement of 5mC alterations in the epigenetic regulation of monocyte exhaustion, we utilized our lab’s previously published *ex vivo* sepsis model in which mouse bone marrow monocytes (BMMCs) are maintained under continuous LPS exposure for five days in the presence of macrophage colony-stimulating factor (M-CSF) (**Fig 1A**) (Pradhan *et al*, 2021). Given that differences in 5mC have been previously implicated in monocyte subtype acquisition, after 5 days of culture, LPS-treated monocytes were sorted by FACS into classical (CD11b^+^;Ly6C^high^), non-classical (CD11b^+^;Ly6C^low^), and intermediate (CD11b^+^;Ly6C^int^) populations; by contrast, PBS-control conditions largely results in a monoculture of Ly6C^low^ non-classical monocytes, which were used as a reference control (Zawada *et al*, 2016). Following bisulfite (BS) conversion, DNA harvested from each subtype population was analyzed using an Infinium Mouse Methylation BeadChip array, which reports on DNA methylation at 285,000 sites across the mouse genome with focused coverage at CpG islands, gene promoters, enhancers, and additional sites of known regulatory significance (Zhou et al, 2022).

**Figure 1.**
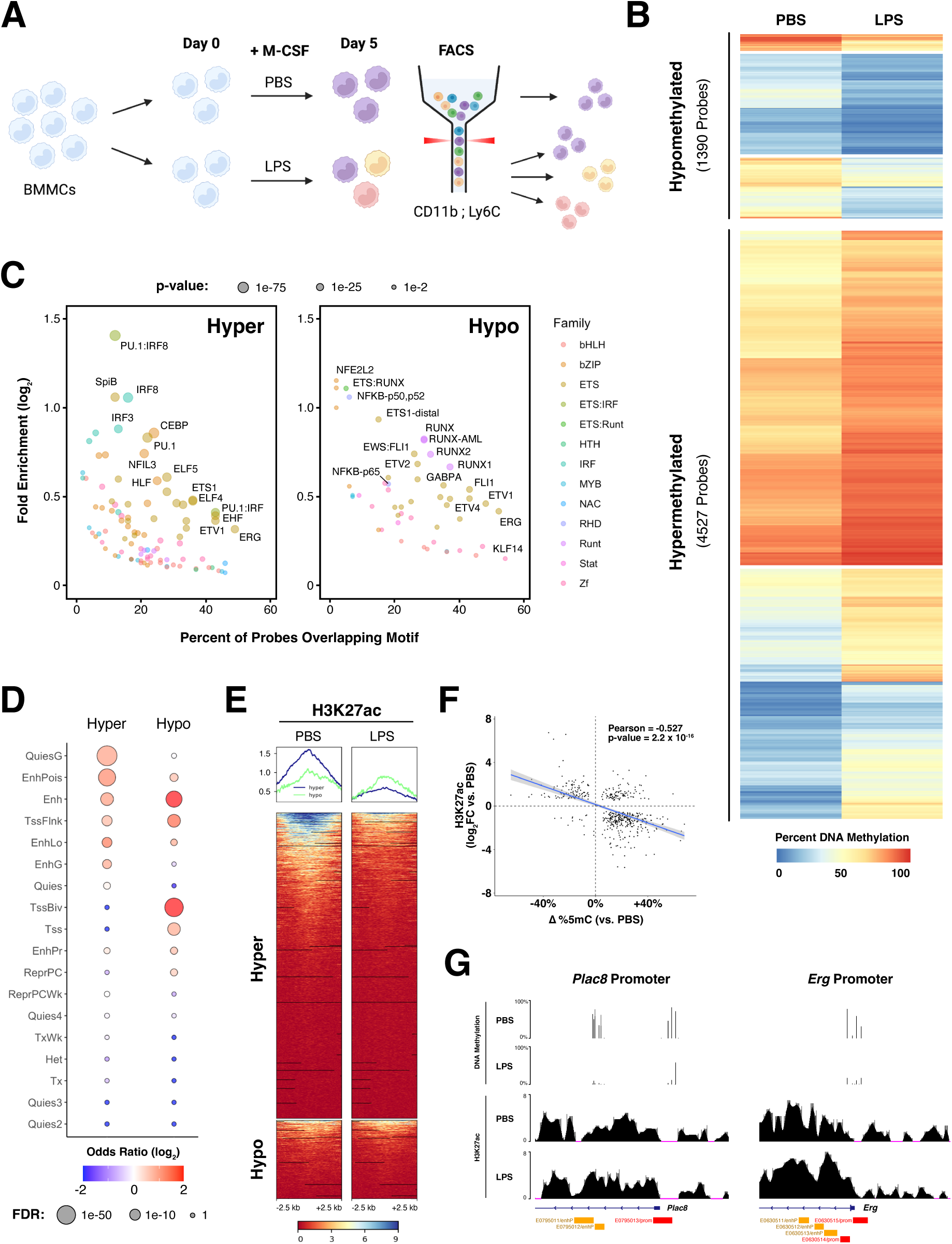
Exhausted monocytes undergo extensive DNA methylation reprogramming in response to repetitive LPS stimulation. **(A)** Schematic for monocyte exhaustion experimental paradigm. Bone marrow monocytes (BMMCs) are cultured under control (PBS) or repetitive LPS stimulation for 5 days in the presence of macrophage colony-stimulating factor (M-CSF). Cells are then sorted into CD111b^+^;Ly6C^low^ (non-classical, purple), CD11b^+^;Ly6C^int^ (intermediate, yellow), or CD11b^+^;Ly6C^high^ (classical, red) pools and analyzed for changes in DNA methylation. **(B)** Heatmap of average DNA methylation at differentially methylated CpG probes following 5 days of PBS control or repetitive LPS stimulation (≥ 5% difference in % 5mC vs. PBS control; FDR < 10%; n = 3 for each monocyte subtype, which was used as an analysis covariate). Rows represent individual probe CpG sites. **(C)** HOMER transcription factor (TF) binding motif analysis for hyper and hypo DMRs (+/- 250 bp non-overlapping windows; bubbles colored by TF family). **(D)** Enrichment of hyper and hypo DMRs in distinct chromatin states using chromHMM. **(E)** Heatmap of H3K27ac enrichment (Data ref: Naler et al., 2022; GEO: GSE168190) at hyper and hypo DMRs in PBS or LPS-treated BMMCs. Metaplots above each heatmap indicated average H3K27ac signal at hyper (blue) and hypo (green) DMRs in each condition (+/- 2.5 kb non-overlapping windows; H3K27ac signal normalized to input control). **(F)** Correlation plot of change in percent DNA methylation vs. log_2_ fold-change in H3K27ac for DMRs with overlapping differential H3K27ac peaks. Simple linear regression was performed, with the line of best fit (blue) and 95% confidence interval (gray highlighted region) indicated. **(G)** UCSC browser track views of MiSeq-validated DMRs at the *Plac8* (chr5:100,570,157-100,573,097) and *Erg* (chr16:95,457,684-95,460,943) promoter regions for BMMCs under control PBS or repetitive LPS stimulation. DNA methylation values represent the average of three biological replicates; values in CD11b^+^;Ly6C^high^ classical monocytes were selected as representative of the LPS condition. ENCODE annotated promoters (red) and enhancers (orange) are depicted in the bottom track.

Global DNA methylation levels were unaffected by LPS stimulation or monocyte subtype, with a median of approximately 85% 5mC observed across all conditions (**Fig EV1A**). Surprisingly, despite their functionally distinct roles in regulating either pro- or anti-inflammatory activity, all three LPS-stimulated monocyte subtypes exhibited near identical changes in 5mC relative to PBS control, suggesting a common suite of differentially methylated regions (DMRs) in exhausted monocytes that underscores their impaired state (**Fig EV1B**). Based on this observation, differential modeling was reperformed using monocyte subtype as a covariate, resulting in the identification of 4,527 hypermethylated (hyper) and 1,390 hypomethylated (hypo) loci (**Fig 1B; Table EV1**). Utilizing sesame’s KnowYourCG (KYCG) probe annotation function, we found that DMRs were heavily enriched at enhancer sites (40% of hyper probes; 32% of hypo) (**Table EV2**) (Zhou *et al*, 2018). Notably, enriched tissue signatures among hyper probes included “monocyte hypomethylated”, while hypo probes included “hematopoietic cell hypomethylated”, suggestive of the emergency hematopoiesis observed in exhausted monocytes (Pradhan *et al*, 2021). Similarly, gene ontology (GO) enrichment of nearest linked genes identified several terms associated with sepsis progression, including “TNF signaling pathway”, “lymphadenopathy”, and “thrombocytopenia” among hypo DMRs and “regulation of actin cytoskeleton” and “IL-3 signaling pathway” among hyper DMRs (**Table EV3**). Interestingly, hypo and hyper DMR-linked genes both demonstrated an enrichment for terms affiliated with G protein signaling, including “GTPase regulator activity”, “CDC42 GTPase cycle”, “RHO GTPase cycle”, and “Ras signaling pathway”, consistent with the known role of these pathways in sepsis-associated dysfunction (Lee et al, 2013b; Hahmeyer & da Silva□Santos, 2021).

Next, we performed HOMER motif analysis to identify transcription factors that may be affected by the altered DNA methylome in exhausted monocytes (**Fig 1C**). Enriched transcription factors within a +/-250bp window of hyper DMRs included numerous members of the ETS, IRF, and CEBP families, which are known to play critical roles in monocyte development and have been implicated in monocyte dysregulation in sepsis and severe Covid-19 infection (Roy *et al*, 2021; Reyes *et al*, 2020; Schulte-Schrepping *et al*, 2020; Kurotaki *et al*, 2013). Interestingly, the PU.1:IRF8 dimer was the most enriched binding motif among hyper DMRs; PU.1 has been shown to modulate DNA methylation levels by recruiting either DNA methyltransferase 3B (DNMT3B) or the DNA demethylation-promoting enzyme TET2, while IRF8 is thought to regulate H3K4me1 levels at enhancers (de la Rica *et al*, 2013; Dekkers *et al*, 2019; Kurotaki *et al*, 2013). In addition to ETS family members, hypo DMRs were heavily enriched for transcription factors involved in RUNX and NF-κB signaling. Whereas NF-κB is known for its role in monocyte tolerogenesis, RUNX proteins are distinguished by their involvement in hematopoiesis (Yan *et al*, 2012; Voon *et al*, 2015). Similar results were obtained using sesame’s KYCG function, which queries transcription factor motifs directly overlapping DMR probes and identified histone reader and Wnt signaling factor MLLT3 as significantly enriched among hyper DMRs (**Fig EV1C**) (Kabra & Bushweller, 2022; Calvanese et al, 2019; Voon et al, 2015).

In agreement with the KYCG probe enrichment analysis, chromatin state discovery and characterization (chromHMM) identified enhancers as heavily enriched at both hyper and hypo DMRs (**Fig 1D**). To better understand how altered DNA methylation at these sites might impact their regulatory function, we utilized our lab’s previously published ChIP-seq data for active histone mark H3K27ac in PBS control or LPS exhausted cultured BMMCs (**Fig 1E**) (Data ref: Naler et al, 2022). Among hyper DMRs, H3K27ac levels were shown to be significantly decreased in LPS-treated cells, consistent with their inactivation in exhausted monocytes. By contrast, H3K27ac levels at hypo DMRs was largely unaffected by LPS stimulation; this pattern is more consistent with the epigenetic priming model for DNA methylation, whereby reduced 5mC levels poises gene enhancers and promoters for future activation in a cell state- or monocyte subtype-dependent manner (Vento-Tormo *et al*, 2016; Lee *et al*, 2021; Dekkers *et al*, 2019; Schmidl *et al*, 2014). Notably, for differential H3K27ac peaks overlapping DMRs, we observed a strong negative correlation between H3K27ac and 5mC levels, further supporting a role for altered DNA methylation in regulating enhancer activity in exhausted monocytes (**Fig 1F**).

Finally, we performed BisPCR^2^ MiSeq to validate 5mC changes observed at immunologically significant sites. Excepting the *Cd80* promoter, significantly altered DNA methylation levels were observed for all tested regions (**Fig EV1D**). These sites include promoters and enhancers for transcription factors known to regulate monocyte behavior (*Foxp1*, *Klf4*, *Runx3*, *Erg*, *Cebpa/g*), cytokines (*Ltb*, *Spn*), and immunomodulatory cell surface receptors (*Pag1, Treml2*, *Trem2*). Remarkably, the strongest DMR observed in exhausted monocytes was hypomethylation in the promoter and intronic enhancer regions of *Plac8* (**Fig 1G**). *Plac8* upregulation has been identified as a top marker for monocyte dysregulation in scRNA-seq datasets collected from both sepsis and severe Covid-19 patients, and its role as an autophagy-linked plasma and lysosomal membrane protein is believed to be essential for SARS-Cov2 infection (Reyes *et al*, 2020; Schulte-Schrepping *et al*, 2020; Ugalde *et al*, 2022; Tse *et al*, 2022). Taken together, the extent of differential methylation at so many sites of immune significance highlights the critical role of 5mC as a regulator of monocyte exhaustion.

### DNA methylation changes are correlated with broad transcriptomic reprogramming in exhausted monocytes

To test the effect of altered DNA methylation on gene expression in exhausted monocytes, we reanalyzed our lab’s previously published scRNA-seq dataset collected from cultured BMMCs under PBS control or repetitive LPS stimulation (Data ref: Pradhan et al, 2021). In agreement with previous scRNA-seq studies, we identified four major monocyte subtypes: Ly6C^high^ classical (C), Ly6C^low^ non-classical (NC), Ly6C^int^ intermediate (Int), and neutrophil-like (Neu) (**Fig2A-B; Fig EV2A**) (Villani *et al*, 2017; Wolf *et al*, 2019; Ikeda *et al*, 2023). Clusters were further subdivided based on the expression of major proliferation markers *Cdc45*, *Cdc6*, *Tpx2*, and *Top2a* (Prolif), as well as based on their enrichment in exhausted BMMC samples over PBS control (Exh), resulting in four exhausted monocyte clusters (ExhC Prolif, ExhC, ExhNC, and ExhNeu) and five control clusters (NC, Int Prolif, Int, Neu Prolif, and Neu). As expected from our flow cytometry experiments, the LPS-stimulated condition was dominated by Ly6C^high^ ExhC cells, while the PBS control condition consisted largely of Ly6C^low^ NC cells (Pradhan *et al*, 2021). Given the poorly characterized role of neutrophil-like monocytes in sepsis progression, we elected to focus on clusters ExhC Prolif, ExhC, and ExhNC for further analysis.

First, DMRs were matched to their nearest gene neighbor, and the difference in DNA methylation observed in exhausted monocytes was correlated with altered gene expression relative to expression in PBS control NC cells (**Fig 2C-D**). As expected, there was a strong correlation between increasing DNA methylation levels and decreased expression, consistent with the known role for DNA methylation in the silencing of gene promoters and enhancers. This effect was typified by *Plac8* and *Socs3*, which exhibit strong hypomethylation of their affiliate regulatory elements concomitant with their increased expression in ExhC cells, while the inverse was observed for *Trem2* and *Cd36*, where hypermethylation correlated with their diminished expression in exhausted monocytes. Interestingly, we also observed altered expression of 5mC writers and erasers in exhausted monocytes; *Dnmt3a* and *Tet3* expression was more strongly enriched in control NC cells, correlating with hypermethylation of their affiliate regulatory elements in exhausted monocytes (**Fig 2E, Table EV1**).

**Figure 2.**
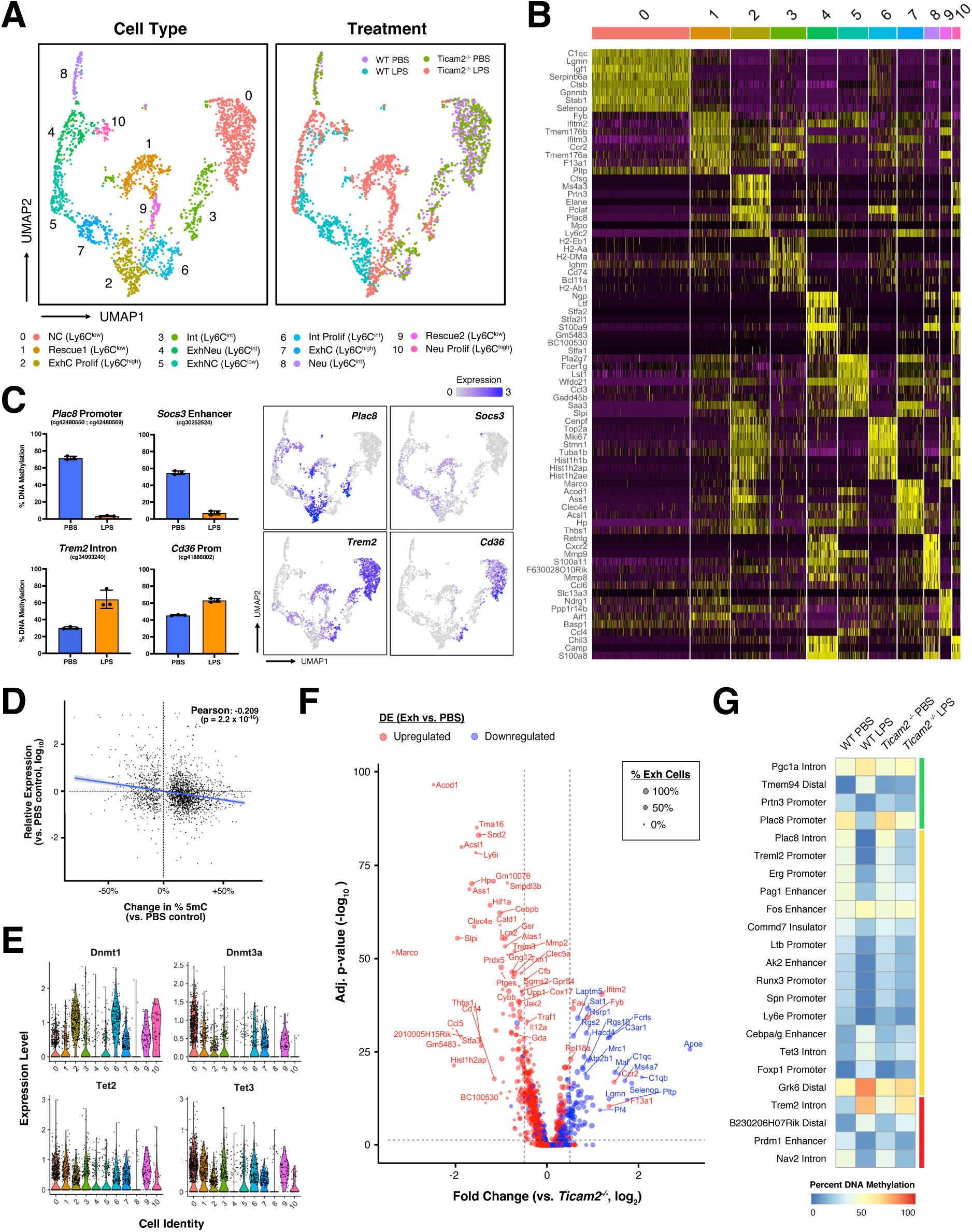
DNA methylation changes correlate with altered gene expression in exhausted monocytes. **(A)** UMAP visualization of scRNA-seq profiles of WT and *Ticam2^-/-^* BMMCs following PBS control or repetitive LPS stimulation (Data ref: Pradhan et al., 2021; GEO: GSE182355). Cells are colored according to monocyte subtype, phenotype, and treatment (NC: non-classical, C: classical, Int: intermediate, Exh: exhausted, Neu: neutrophil-like, Prolif: proliferating). **(B)** Heatmap of intersectional gene expression for the top 8 markers of each cell cluster. **(C)** Promoter (*Plac8*, *Trem2*, *Cd36*) and enhancer (*Socs3*) DMRs for immunologically significant genes in WT PBS control or Ly6C^high^ LPS-treated BMMCs with affiliate UMAP feature plots demonstrating expression levels across cell clusters. Illumina BeadChip probe IDs are indicated for each DMR. **(D)** Correlation plot of change in percent DNA methylation vs. log_10_ fold- change in expression for the nearest matched gene between WT PBS control and Ly6C^high^ LPS-treated BMMCs. Simple linear regression was performed, with the line of best fit (blue) and 95% confidence interval (gray highlighted region) indicated. **(E)** Gene expression levels across cell clusters for known regulators of DNA methylation. **(F)** Volcano plot depicting log_2_ fold-change in exhaustion gene expression between LPS- treated WT (clusters 2, 5, and 7) and *Ticam2^-/-^* (clusters 1, 2, and 5) BMMCs. Genes are colored by the observed differential expression pattern in WT exhausted cells vs. PBS control (cluster 0), while dot sizes are scaled based on the proportion of WT exhausted cells expressing the indicated gene. Dotted lines indicate the fold change (1.5x) and adjusted p-value (< 0.05) cut-offs for differentially expressed genes. **(G)** Heatmap of average DNA methylation levels at MiSeq-validated DMRs following 5 days of PBS control or repetitive LPS stimulation in WT or *Ticam2^-/-^*BMMCs. Colored bars indicate full (green), partial (orange), or failed (red) rescue status of DMRs in *Ticam2^-/-^* LPS- treated cells (n = 3-9 per condition).

Previous research from our group has implicated TLR4 signaling component TICAM2 as a major upstream regulator of myeloid exhaustion (Lin *et al*, 2020; Pradhan *et al*, 2021). To better understand the role of TICAM2 in establishing the altered 5mC landscape of exhausted monocytes, we integrated scRNA-seq data from *Ticam2^-/-^* BMMCs under PBS control or repetitive LPS stimulation into our single-cell analysis. Whereas control *Ticam2^-/-^* BMMCs largely clustered with their wild-type (WT) counterparts, *Ticam2^-/-^* BMMCs cultured under repetitive LPS stimulation exhibited distinct clustering within Exh populations relative to WT cells (**Fig 2A**). Notably, *Ticam2^-/-^* samples were depleted of ExhC cells, instead forming two unique Ly6C^low^ populations (Rescue 1 and 2) that more closely resembled PBS control NC cells. To evaluate the degree of rescue in transcriptional reprogramming of *Ticam2^-/-^* BMMCs, we determined the fold change expression of WT LPS-treated cells (ExhC Prolif, ExhC, and ExhNC) relative to *Ticam2^-/-^* (Rescue 1, 2, and Int), which identified 76 upregulated and 53 downregulated genes whose expression was significantly rescued in *Ticam2^-/-^* BMMCs (**Fig 2F**). Consistent with this altered transcriptional profile, BisPCR^2^ sequencing of representative exhaustion DMRs revealed at least a partial rescue of DNA methylation levels in 19/23 tested regions (**Fig 2G, Fig EV2B**). Taken together, these data implicate TICAM2 as a major upstream regulator of monocyte exhaustion at both the levels of transcriptomic and 5mC reprogramming.

Finally, to directly assay the influence of altered DNA methylation on transcriptomic changes observed in exhausted monocytes, we cultured BMMCs in the presence of DNMT inhibitor 5-azacytidine (5-aza) during PBS control or repetitive LPS stimulation. 5-aza was well tolerated by BMMCs at concentrations up to 250 nM, with successful DNMT inhibition confirmed by BisPCR^2^ sequencing (**Fig 3A**). Whereas 5-aza treatment had a minimal effect on gene expression in PBS control cells, transcriptional suppression of numerous exhaustion genes was noted in BMMCs under repetitive LPS stimulation, including *Plac8*, *Socs3*, *Runx3*, and *Cd38* (**Fig 3B**). Importantly, the expression of canonical tolerance gene *Acod1* was unaffected by 5-aza treatment, indicating the IM tolerance apparatus remained largely intact in 5-aza-treated cells (Li *et al*, 2013). Intriguingly, expression of exhaustion marker *Mpo* and was strongly upregulated by 5-aza treatment, suggesting that hypermethylation observed in its promoter region in LPS-treated BMMCs may be critical for its limited activation (**Table EV1**). We also evaluated 5-aza rescue by flow cytometry for known exhaustion markers. In agreement with the limited differences in DNA methylation among exhausted monocyte subtypes, 5-aza treatment had a negligible impact on the relative levels of C, NC, and Int monocytes (**Fig 3C**). While cell surface levels of tolerance marker PD-L1 were similarly unaffected by 5-aza treatment, CD38 levels were greatly diminished, and CD86 nearly reached levels observed in PBS control cells (**Fig 3D**) (Avendanõ-Ortiz *et al*, 2018a). Given that the majority of DNA methylation changes observed exhausted BMMCs were from gain of 5mC, it is likely that pharmacological inhibition of DNMT limits the extent of exhaustion phenotype acquisition in LPS-treated cells, supporting the critical involvement of DNA methylation in this process.

**Figure 3.**
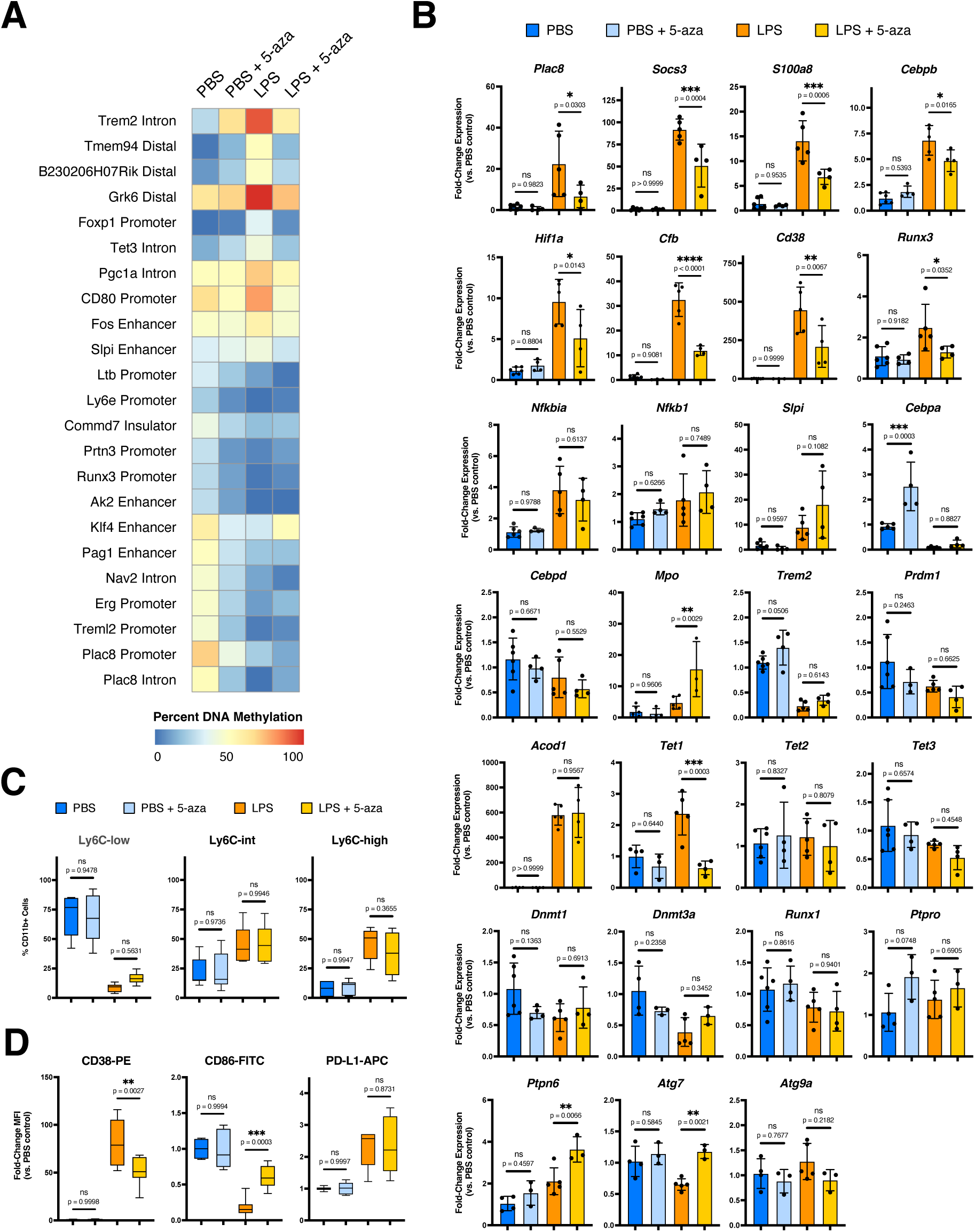
DNA methyltransferase inhibition during repetitive LPS stimulation of BMMCs alleviates the exhaustion phenotype. **(A)** Heatmap of average DNA methylation levels at MiSeq-validated DMRs in BMMCs following 5 days of PBS control or repetitive LPS stimulation in the presence of 250 nM 5-azacytidine (5-aza). **(B)** qRT- PCR for key exhaustion gene expression relative to PBS control. Expression levels were normalized to the geometric mean of *Ube2l3*, *Oaz1*, and *Nktr*. Data points represent independent experiments, with mean expression +/- STD indicated (n = 3-6; one-way ANOVA with Sidak’s multiple comparisons test; **** p-adj. < 0.0001; *** < 0.001; ** < 0.01; * < 0.05; ns not significant). **(C)** Population statistics for non-classical (Ly6C^low^), intermediate (Ly6C^int^), and classical (Ly6C^high^) monocyte subtypes in cultured BMMCs (n = 6-7 ; one-way ANOVA with Sidak’s multiple comparisons test). **(D)** Flow cytometry mean fluorescence intensity (MFI) for exhaustion markers relative to PBS control (n = 6-7; one-way ANOVA with Sidak’s multiple comparisons test).

### Wnt signaling antagonizes DNA methylation changes and transcriptomic reprogramming in exhausted monocytes

We next performed SCENIC analysis to more precisely identify those regulatory circuits involved in the transcriptomic and 5mC reprogramming of exhausted monocytes (Aibar et al, 2017). In total, we identified 63 high-confidence regulons spanning all scRNA-seq identified BMMC clusters (**Fig 4A**). Notably, several transcription factors previously implicated in monocyte dysregulation of sepsis and severe Covid-19 patients were significantly enriched in our exhausted cell clusters, including HIF1A, CEBPB, CEBPD, and NF-KB2 (**Fig 4B**) (Reyes *et al*, 2020; Schulte-Schrepping *et al*, 2020; Avendanõ-Ortiz *et al*, 2018b; Kang *et al*, 2016). Consistent with RUNX3 binding motif enrichment at hypo DMRs in LPS-treated cells, we also observed distinct RUNX3 activity in ExhC Prolif cells, again supporting a role for RUNX3-driven emergency hematopoiesis in this population (**Fig 1C**). Interestingly, despite observing similar DNA methylation patterns in exhausted monocytes regardless of Ly6C cell surface levels, regulons in the four Exh clusters were largely non-overlapping. As with our H3K27ac ChIP-seq analysis, this suggests DNA demethylation in exhausted monocytes functions to prime the genome for future gene activation in a subtype-specific manner.

**Figure 4.**
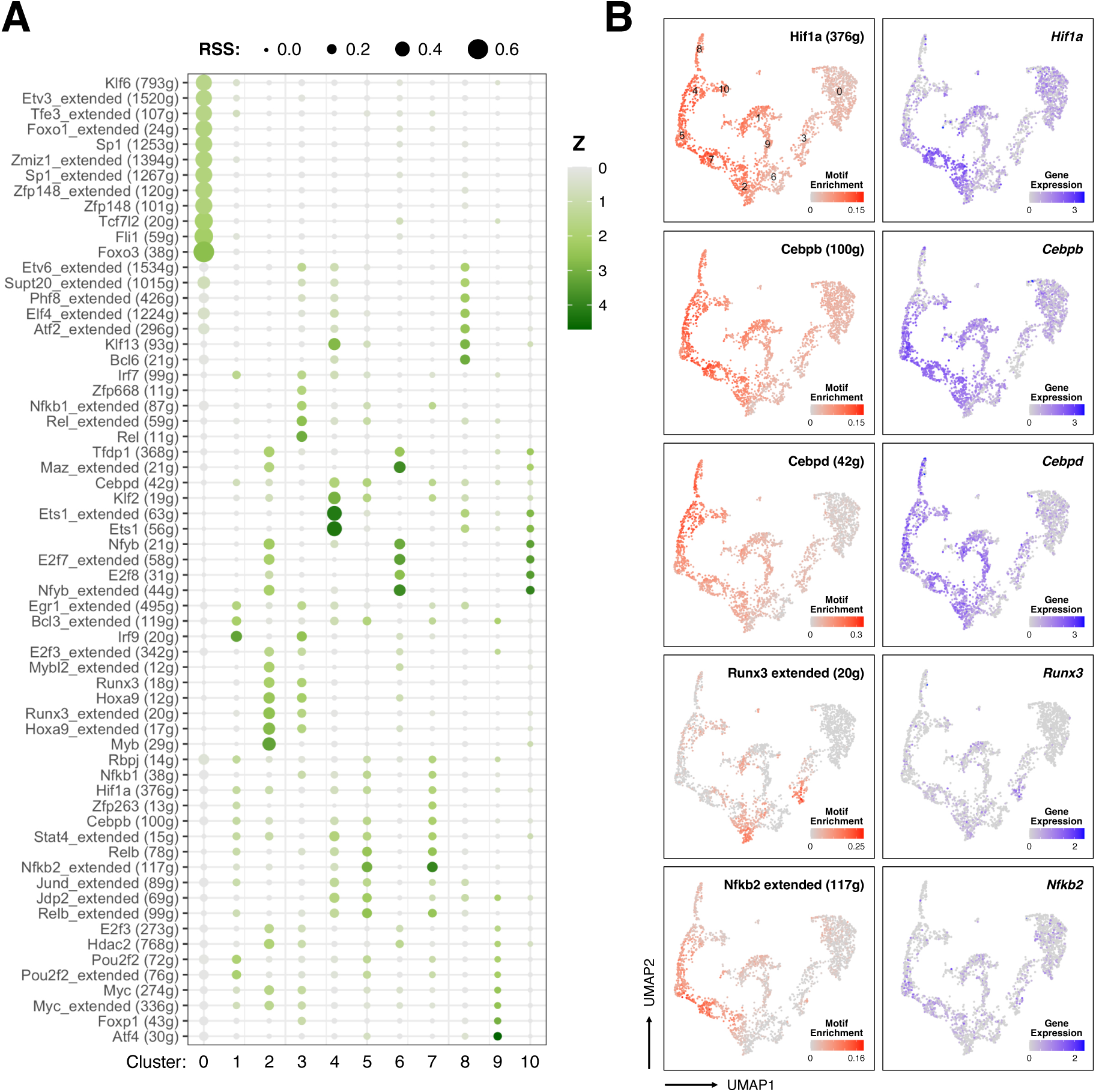
SCENIC analysis of exhausted monocyte scRNA-seq data. **(A)** Dot plot for SCENIC transcription factor motif enrichment in WT and *Ticam2^-/-^* BMMCs following PBS control or repetitive LPS stimulation. Dot sizes represent a given motifs regulon specificity score (RSS) for the indicated cell cluster. Numbers represent different cell clusters, as outlined in Figure 2A (RSS cut-off = 0.1; Z-threshold = 1.5). **(B)** scRNA-seq UMAP feature plots for key exhaustion transcription factor SCENIC motif enrichment and gene expression in cultured BMMCs.

Among regulons enriched in PBS control cells, we noted several involved in anti-inflammatory, pro-resolving monocyte behavior, including FOXO3, FLI1, ETV3 (**Fig 4A**) (El Kasmi *et al*, 2007; Bujor *et al*, 2020; Lee *et al*, 2013a). Of particular interest was TCF7L2, a downstream effector of canonical Wnt signaling most well-known for its strong genetic association with type 2 diabetes risk (**Fig 5A**) (Del Bosque-Plata *et al*, 2021). Given the poorly characterized role of TCF7L2 and Wnt signaling in monocyte biology, we were interested testing the effect of Wnt signaling activation during monocyte exhaustion. PBS control and LPS-stimulated BMMCs were treated with 1 or 5 µM Wnt Agonist 1 starting at day 2 of cell culture. Strikingly, Wnt signaling activation had a profound effect on monocyte exhaustion at the transcriptional level, limiting the expression of exhaustion genes such as *Plac8*, *Cd38*, and *Cfb* in addition to Exh regulon members *Hif1a* and *Runx3* (**Fig 5B**). Despite a modest decrease in *Nfkb1* expression in Wnt agonist-treated cells under LPS stimulation, we noted that *Acod1* expression remained elevated, suggesting tolerance is unimpacted by increased Wnt signaling (**Fig 5B, EV3A**).

**Figure 5.**
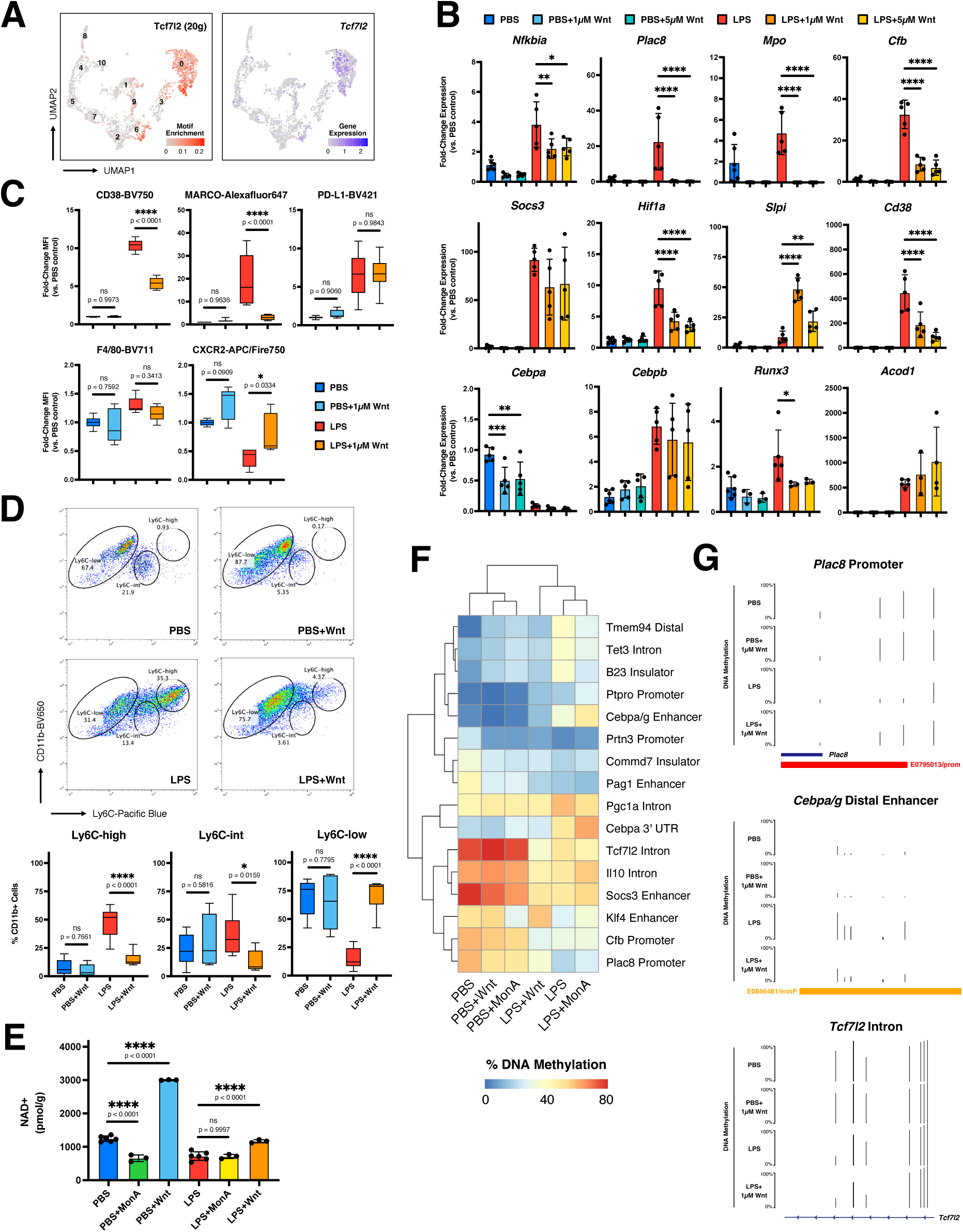
Wnt signaling activation limits innate immune exhaustion phenotype and DNA methylation alterations observed in BMMCs under repetitive LPS stimulation. **(A)** scRNA-seq UMAP feature plots for transcription factor Tcf7l2 motif enrichment and gene expression in WT and *Ticam2^-/-^* BMMCs following PBS control or repetitive LPS stimulation. Numbers indicate different cell clusters, as outlined in Figure 2A. **(B)** qRT-PCR for key exhaustion gene expression in BMMCs under PBS control or repetitive LPS stimulation in the presence of different concentrations of Wnt Agonist 1. Expression levels were normalized to the geometric mean of *Ube2l3*, *Oaz1*, and *Nktr*. Data points represent independent experiments, with mean expression +/- STD indicated (n = 3-6; one-way ANOVA with Sidak’s multiple comparisons test; **** p-adj. < 0.0001; *** < 0.001; ** < 0.01; * < 0.05; ns not significant; differences are not significant unless otherwise specified; see Appendix for exact p-values). **(C)** Flow cytometry mean fluorescence intensity (MFI) for exhaustion (CD38, MARCO, PD-L1, CXCR2) and macrophage (F4/80) markers relative to PBS control. Box plots indicate median MFI values (n = 5-7; one-way ANOVA with Sidak’s multiple comparisons test). **(D)** Gating strategy (top) and population statistics (bottom) for non-classical (Ly6C^low^), intermediate (Ly6C^int^), and classical (Ly6C^high^) monocyte subtypes in cultured BMMCs (n = 6-13 ; one-way ANOVA with Sidak’s multiple comparisons test). **(E)** NAD+ levels in cultured BMMCs normalized to total protein levels (n = 3-6; one-way ANOVA with Sidak’s multiple comparisons test). **(F)** Correlation heatmap for average DNA methylation levels at key exhaustion loci in cultured BMMCs. (n = 3-9 for each condition). **(G)** UCSC browser track views of average DNA methylation at the *Plac8* promoter (chr5:100,572,230-100,572,607) and *Cebpa/g* distal enhancer (chr7:35,064,805-35,065,304), and *Tcf7l2* intron (chr19:55,768,986-55,769,585) regions in cultured BMMCs. ENCODE annotated promoters (red) and enhancers (orange) are depicted in the bottom track.

We next performed flow cytometry to further characterize the effect of Wnt Agonist 1 treatment on monocyte exhaustion. In agreement with our qRT-PCR results, cell surface levels of exhaustion markers CD38 and MARCO were significantly depleted following Wnt Agonist 1 treatment, while PD-L1 levels remained unaffected (**Fig 5C**). We also observed an increase in cell surface levels of the IL8-responsive G-protein-coupled receptor CXCR2, which is normally reduced in exhausted monocytes. Although Wnt signaling has been reported to block monocyte-to-macrophage differentiation, we confirmed that macrophage marker F4/80 cell surface levels were negligibly impacted by our low-dose Wnt Agonist 1 intervention. In contrast to 5-aza treatment, we observed a substantial repartitioning of monocyte subtypes in Wnt Agonist 1-treated BMMCs under LPS stimulation (**Fig 5D**). The proportion of Ly6C^low^ NC monocytes increased to PBS control levels, while Ly6C^int^ and Ly6C^high^ monocytes were significantly depleted. In concert with the observed gene expression changes, these results suggests that Wnt signaling activation promotes an anti-inflammatory, pro-resolving state in monocytes under repetitive LPS stimulation.

To test the specificity of Wnt-mediated suppression of the exhaustion phenotype, we treated PBS control and LPS-stimulated BMMCs with Wnt signaling antagonist monensin A (MonA) (Tumova *et al*, 2014). In contrast to Wnt Agonist 1 treatment, transcription was largely unimpacted by MonA (**Fig EV3B**), although *Plac8*, *Mpo*, and *Cfb* were still observed to be downregulated in MonA-treated cells under LPS stimulation. Of note, all three of these genes are direct targets of c-MYB, which has enriched regulatory activity in ExhC cluster cells (**Fig EV3C**) (Zhao *et al*, 2011). Given that MonA was identified as a potent chemical inhibitor of c-MYB, it is probable that MonA-mediated repression of these genes proceeds through decreased c-MYB activity (Yusenko *et al*, 2020). Flow cytometry analysis of MonA-treated cells further confirmed that cell surface levels of CD38 and CD86 were unaffected by Wnt inhibition, and that PD-L1 levels actually increased in LPS-treated cells (**Fig EV3D**). Surprisingly, while the proportion of pro-inflammatory Ly6C^high^ monocytes was increased in MonA-treated cells under LPS stimulation, PBS control cells were completely depleted of Ly6C^high^ and Ly6C^int^ cells (**Fig EV3E**). Such an effect mirrors the subtype repartitioning observed in Wnt Agonist 1-treated BMMCs under LPS stimulation, and suggests a context-dependent role for Wnt signaling in regulating monocyte behavior.

We and others have previously reported CD38-mediated NAD^+^ depletion in sustained pathogenic inflammation and monocyte exhaustion (Piedra-Quintero *et al*, 2020; Joe *et al*, 2020; Hong *et al*, 2018; Galley, 2011; Pradhan *et al*, 2021). To determine the impact of Wnt signaling on mitochondrial dysfunction in exhausted monocytes, we measured NAD^+^ levels in LPS-treated BMMCs during Wnt signaling intervention. While NAD^+^ levels remained depleted in LPS-treated cells following MonA treatment, NAD^+^ levels in PBS control cells was significantly diminished by MonA, consistent with the drug’s known effect as a driver of mitochondrial disruption and oxidative stress (**Fig 5E**) (Charvat & Arrizabalaga, 2016). By contrast, Wnt Agonist 1 treatment led to a substantial increase in NAD^+^ in both PBS control and LPS-treated BMMCs, with NAD^+^ in LPS-treated cells rising to control levels. To further characterize the metabolic impact of Wnt signaling during monocyte exhaustion, we measured autophagy gene *Atg7* expression, which is significantly diminished in exhausted monocytes. Whereas *Atg7* remains repressed in MonA-treated cells, its expression was fully rescued in Wnt Agonist 1-treated cells (**Fig EV3A-B**). Promotion of Wnt signaling thus seems to have a positive impact on metabolic stress in BMMCs under repetitive LPS stimulation.

Finally, to determine the effect of Wnt signaling on 5mC reprogramming in exhausted monocytes, we performed BisPCR^2^ sequencing for select exhaustion DMRs following MonA or Wnt Agonist 1 treatment. While the effect of either treatment on DNA methylation in PBS control cells was fairly limited, Wnt Agonist 1 treatment rescued 5mC levels at several DMRs in LPS-treated cells, including the *Cebpa/g* enhancer, *Plac8* promoter, and *Cebpa* 3’UTR (**Fig 5F-G**). Interestingly, whereas Wnt Agonist 1 treatment rescued 5mC levels at the *Klf4* enhancer in LPS-treated cells, MonA treatment drove hypomethylation of this element in PBS control cells to levels normally observed in exhausted monocytes. It is also noteworthy that *Tcf7l2* itself exhibits hypomethylation of an unannotated intronic region in exhausted monocytes, and that this effect is slightly exacerbated by Wnt Agonist 1 treatment (**Fig 5G**). Taken together, these results demonstrate the Wnt signaling intervention in exhausted monocytes not only has the capacity to reverse transcriptional and metabolic dysregulation in these cells, but also to impede the establishment of 5mC memory states at critical regulatory genes.

### TDM-mediated immune training inhibits 5mC reprogramming in exhausted monocytes

The interplay between training and tolerance mechanisms remains an active area of research. Previous work has demonstrated that treatment of tolerized monocytes with immune training compound β-glucan can restore H3K27ac levels at promoters and distal enhancers and rescue the expression of roughly 60% of tolerized genes (Novakovic *et al*, 2016). However, the effect of immune training on DNA methylation changes during LPS stimulation remains unexplored, particularly in the context of monocyte exhaustion.

Trehalose 6,6’-dimycolate (TDM), an abundant glycolipid component in the mycobacterial cell wall, was one of the earliest identified immunostimulatory molecules capable of driving immune training in treated animals (Decout *et al*, 2017; Madonna *et al*, 1989a). To test the effect of immune training on 5mC reprogramming in exhausted monocytes, we cultured BMMCs under repetitive LPS stimulation for five days in the presence of TDM. Whereas global DNA methylation levels relative to PBS control cells were unaffected by LPS or TDM treatment, there was a substantial reduction in hypermethylated regions in LPS-treated cells stimulated with TDM (10,533 LPS vs. 3,037 LPS+TDM) (**Fig 6A-B, Fig EV4A**, **Table EV4)**. DMRs observed in LPS+TDM-treated cells largely overlapped those observed in BMMCs treated with LPS alone (88% of hyper DMRs; 74% of hypo), although there were a substantial number of hypo DMRs unique to LPS+TDM treatment. Principal components analysis of 5mC patterns supported a clear separation of LPS- and LPS+TDM-treated BMMCs, with component 2 being driven primarily by unique differences observed in TDM-treated cells (**Fig 6C**). To more effectively quantify the extent of TDM-mediated rescue of 5mC reprogramming in exhausted monocytes, we reperformed differential 5mC analysis using BMMCs treated with LPS alone as the reference control (**Fig 6D**). In total, 2,636 hyper and 39 hypo exhaustion DMRs were significantly rescued by TDM treatment (> °5% Δ5mC; adjusted p-value < 0.05). This effect was clearly observed in several sites of regulatory (*Pim1*, *Foxp1*) and immune (*Il12rb1*, *Nfatc2*, *Socs3*, *CD83*) importance, supporting improved immunogenicity in TDM-trained cells (**Fig 6E**). Additionally, we also identified 133 significant DMRs unique to the LPS+TDM treatment. Of particular note was a strong reduction in DNA methylation at an intronic enhancer for H3K79 methyltransferase gene *Dot1l*; given the important role of histone modifications in immune training, DNA demethylation may facilitate this process through the transcriptional activation *Dot1l* (Wood et al, 2018).

**Figure 6.**
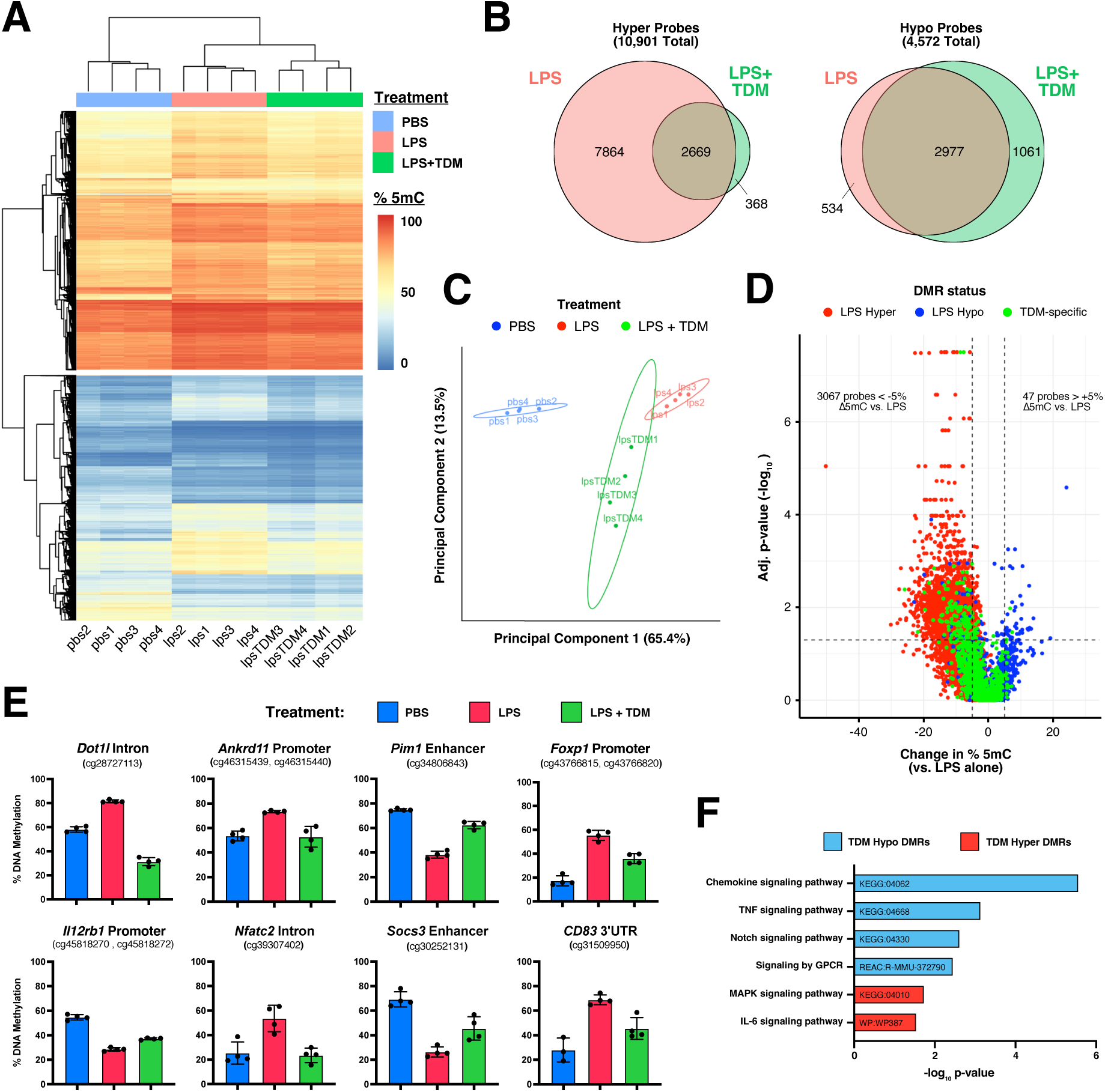
Immune training with TDM limits DNA methylation changes in BMMCs under repetitive LPS stimulation. **(A)** Correlation heatmap for DNA methylation at differentially methylated CpG probes following 5 days of PBS control or repetitive LPS stimulation in the presence or absence of TDM (≥ 5% difference in % 5mC vs. PBS control; FDR < 5%; n = 4 for each treatment). **(B)** Venn diagrams for differentially methylated CpG probes in cells treated with LPS alone (red) or LPS + TDM (green). **(C)** Principal component analysis of differentially methylated CpG probes in PBS control- (blue), LPS- (red), or LPS+TDM-treated (green) BMMCs. Ovals indicate the normal distribution for each treatment. **(D)** Volcano plot for altered DNA methylation at DMRs in LPS+TDM-treated BMMCs versus treatment with LPS alone. Probes are colored based on observed differential methylation patterns in LPS-treated cells relative to PBS control (red: hypermethylated; blue: hypomethylated) or probes that exhibit differential methylation only in LPS+TDM cells (green). Dotted lines indicate the change in DNA methylation (+/- 5%) and adjusted p-value (< 0.05) cut-offs for DMRs. **(E)** Representative DMRs linked to epigenetic modifiers (*Dot1l*, *Ankrd11*), transcriptional regulators (*Pim1*, *Foxp1*), and immunologically significant genes (*Il12rb1*, *Nfatc2*, *Socs3*, *CD83*) in PBS control- (blue), LPS- (red), or LPS+TDM-treated (green) BMMCs. Illumina BeadChip probe IDs are indicated for each DMR. **(F)** Gene ontology (GO) analysis for enriched cell signaling pathways among DMR-linked genes in LPS+TDM- treated BMMCs.

We next annotated DMR probes in LPS+TDM BMMCs using sesame’s KYCG function (**Table EV5**). As with BMMCs under LPS stimulation alone, DMRs were heavily enriched at enhancer sites (43% of hyper probes; 34% of hypo). This effect was confirmed by chromHMM analysis, which demonstrate similar chromatin enrichment of LPS+TDM DMRs regardless of whether the site was unique to the LPS+TDM condition (**Fig EV4B**). Tissue signature enrichment still identified “monocyte hypomethylated” among hyper DMRs and “hematopoietic cell hypomethylated” among hypo DMRs, suggesting that while 5mC reprogramming is diminished in TDM-treated cells, monocyte exhaustion is still apparent at the level of DNA methylation (**Table EV5**). We also identified a similar repertoire of transcription factor binding sites overlapping DMRs in TDM+LPS-treated cells, although MLLT3 enrichment was substantially diminished at hyper DMRs along with CEBP family members CEBPA, CEBPB, and CEBPE (**Fig EV4C**). Given that Wnt signaling was shown to antagonize monocyte exhaustion, general loss of DNA methylation at the binding sites for Wnt factor MLLT3 likely promotes Wnt signaling in LPS+TDM-treated BMMCs and contribute to their improved immune function.

Finally, we performed GO enrichment analysis of DMR-linked genes to pinpoint molecular pathways that may contribute to immune training in TDM-treated cells (**Table EV6**). As with BMMCs treated with LPS alone, we noted significant enrichment of “TNF signaling pathway” and “Signaling by GPCR” among hypo DMRS (**Fig 6F**). However, altered immunogenicity in TDM-treated cells was evident from enrichment of “chemokine signaling” and “Notch signaling” among hypo DMRs and “MAPK signaling” and “IL-6 signaling” among hyper DMRs. In concert with NF-κB, MAPK signaling has been shown to promote HIF-1 activation during LPS stimulation, while a Notch and JAK/STAT3 positive feedback loop is thought to drive downstream IL-6 secretion and PD-L1 expression during LPS stimulation (Hildebrand *et al*, 2018; Ohishi *et al*, 2000; Frede *et al*, 2006). Altered DNA methylation of these immune signaling pathways’ affiliate members is expected to regulate their expression during immune challenge, further highlighting the potential benefits of immune training on monocyte exhaustion.

### In vivo evidence for DNA methylation involvement in monocyte exhaustion and immune memory

Based on our *ex vivo* mouse sepsis model, DNA methylation is significantly altered in bone marrow-derived monocytes under repetitive LPS stimulation. These changes correlate strongly with transcriptional reprogramming in exhausted monocytes, with DMRs identified at key regulatory sites for sepsis genes just such as *Plac8*, *Runx3*, *Klf4*, *Socs3*, and *Foxp1*. To better understand the role of DNA methylation plays in regulating monocyte behavior *in vivo*, we performed cecal slurry injections to induce sepsis in wild-type C57BL/6J mice. Injections were performed at day 0 (d0) and d5 of the experimental paradigm, after which bone marrow samples were collected at d6, d7, and d12 (**Fig 7A**).

**Figure 7.**
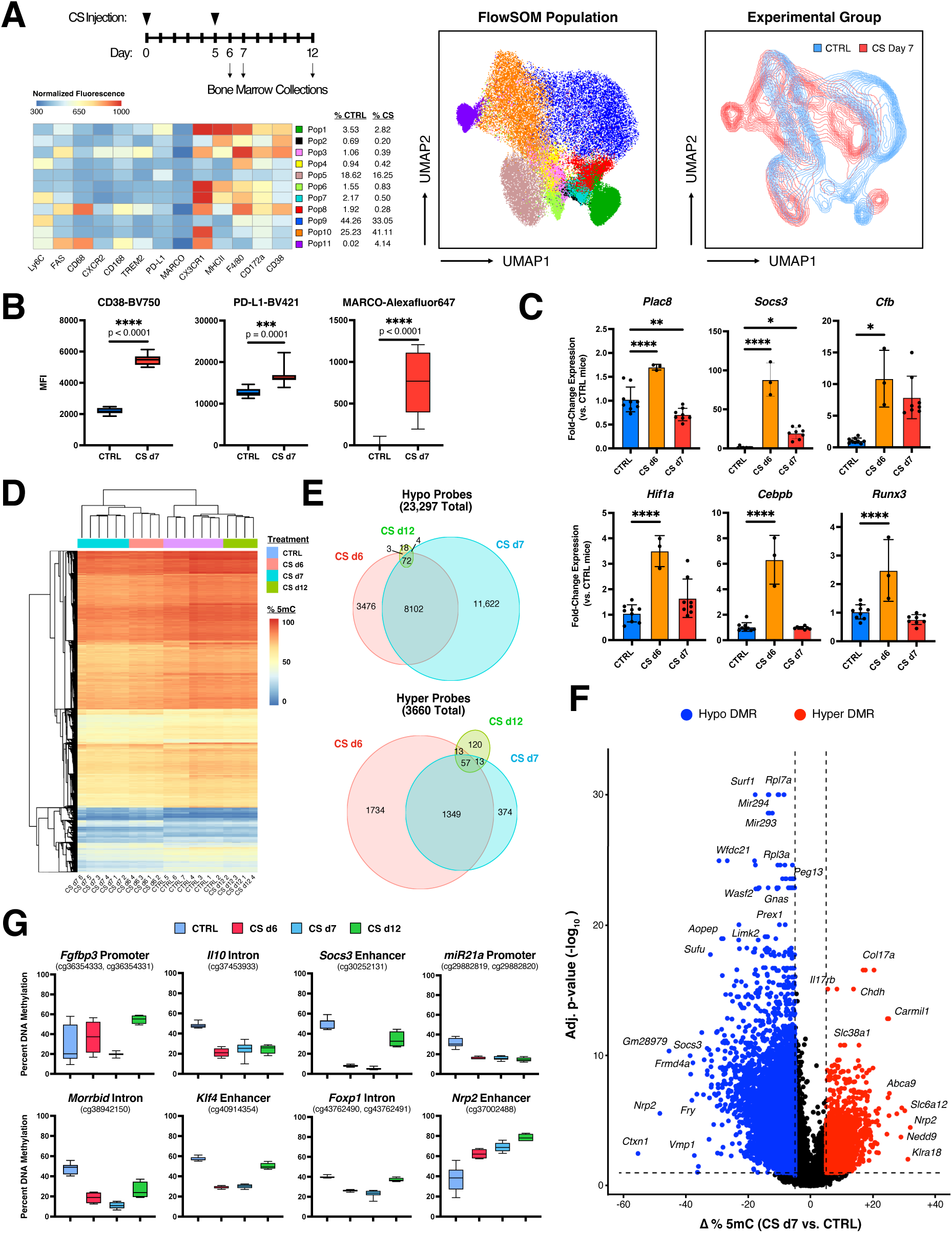
*In vivo* validation of major regulatory circuit and altered DNA methylome in exhausted monocytes. **(A)** Experimental paradigm and FlowSOM analysis of monocytic lineage populations (CD11b^+^; Ly6g-) in bone marrow samples collected from control (CTRL) or cecal slurry (CS)-injected mice at experimental day 7 (d7). Percentages to the right of each FlowSOM population indicate the percentage of cells for each condition clustering in that population. UMAP visualizations were prepared using flow cytometry data for the indicated markers (n = 8 CTRL and 12 CS d7 mice). **(B)** Flow cytometry mean fluorescence intensity (MFI) for exhaustion markers in FlowSOM Population 1. Box plots indicate median MFI values (n = 8-12; one-way ANOVA with Sidak’s multiple comparisons test; **** p-adj. < 0.0001; *** < 0.001; * < 0.05). **(C)** qRT-PCR for key exhaustion gene expression in CTRL and CS d6 or d7 monocytes. Expression levels were normalized to the geometric mean of *Ube2l3*, *Oaz1*, and *Nktr*. Data points represent separate mice, with mean expression +/- STD indicated (n = 3-9; one-way ANOVA with Sidak’s multiple comparisons test; differences are not significant unless otherwise specified; see Appendix for exact p-values). **(D)** Correlation heatmap for DNA methylation at differentially methylated CpG probes in CTRL or CS bone marrow monocytes (significance defined by ≥ 5% difference in % 5mC vs. CTRL; FDR < 10%; n = 7 CTRL, 4 CS d6, 6 CS d7, and 4 CS d12 mice). **(E)** Venn diagrams for differentially methylated CpG probes in CS bone marrow monocytes at experimental days 6, 7, and 12. **(F)** Volcano plot for altered DNA methylation at DMRs in CS d7 bone marrow monocytes versus CTRL. Probes are colored based on observed differential methylation state in CS d7 monocytes (red: hypermethylated; blue: hypomethylated). Dotted lines indicate the change in DNA methylation (+/- 5%) and adjusted p-value (< 0.1) cut-offs for DMRs. Nearest linked genes are indicated for select DMRs. **(G)** Promoter (*Fgfbp3*, *miR21a*, *Cd36*) or enhancer (*Il10, Socs3, Morrbid, Klf4, Foxp1, Nrp2*) DMRs for immunologically significant genes in CTRL or CS monocytes. Illumina BeadChip probe IDs are indicated for each DMR.

Given myeloid lineage diversity in bone marrow-derived cells, we developed a 16-factor flow cytometry panel from our scRNA-seq dataset to more effectively characterize the exhaustion phenotype in bone marrow samples. After gating for non-neutrophil, monocytic lineage cells (Ly6g^-^;CD11b^+^), we performed Flow Self-Organizing Map (FlowSOM) automated cell clustering using phenotypic markers for monocytes/macrophages (Ly6C, MARCO, CD68, CX3CR1, MHCII, F4/80, CD172a), proliferation (CD168), exhaustion (FAS, CD38, TREM2, CXCR2), and tolerance (PD-L1) on bone marrow collected from d7 cecal slurry (CS) or control (CTRL) mice (**Fig7A**). (Van Gassen *et al*, 2015). Major cellular repartitioning in CS samples was noted for nearly every cell cluster identified, with particular depletion of inflammatory populations (Pop) 2, 3, 6, 7, and 8 relative to CTRL samples, likely due to their emigration to sites of infection during sepsis (**Fig EV5A**) (Serbina *et al*, 2008; Cebinelli *et al*, 2021). While no significant difference was observed in the proportion of mature (F4/80^+^) Pop1 macrophages, CS Pop1 cells exhibited a marked exhaustion phenotype of heightened CD38, PD-L1, and MARCO expression (**Fig 7B**). We also noted a near two-fold increase in lowly differentiated Pop10 Ly6C^int^ monocytes in CS bone marrow samples, which is commonly observed in both septic shock and severe Covid-19 infection (**Fig EV5A**) (Mukherjee *et al*, 2015; Gibellini *et al*, 2020; Hortová-Kohoutková *et al*, 2020). Furthermore, we observed a Ly6C^int^ monocyte population (Pop11) unique to CS bone marrow samples with elevated levels of inflammatory and phagocytic markers CX3CR1 and CD68 in addition to high expression of pro-apoptotic TNF-receptor superfamily member FAS, a notable feature of sepsis pathogenesis (Ayala et al, 2003; Wesche et al, 2005).

To validate the exhaustion regulatory apparatus identified in *ex vivo* septic BMMCs, we measured the expression of key exhaustion effector (*Plac8*, *Socs3*, *Cfb*) and regulatory (*Hif1a*, *Cebpb*, *Runx3*) genes in monocytes column purified from d6 and d7 CS bone marrow samples (**Fig 7C**). In agreement with *ex vivo* results, we noted a transient enrichment for *Plac8*, *Hif1a*, *Cebpb*, and *Runx3* expression in CS d6 samples relative to CTRL. Whereas expression of these genes normalized in d7 monocyte samples, *Socs3* and *Cfb* remained elevated, representative of the paradoxical pro-inflammatory (*Cfb*), anti-immune (*Socs3*) behavior of exhausted monocytes.

Next, we analyzed DNA methylation in column purified bone marrow monocyte samples using Infinium Mouse Methylation BeadChip Arrays to identify changes during acute sepsis (CS d6 and d7). Progressive, genome-wide hypomethylation was observed in CS d6 and d7 monocytes relative to CTRL (median % 5mC = 78% in CTRL cells versus 76% and 75% in CS d6 and d7, respectively), consistent with previous reports that monocyte differentiation and activation is characterized by loss of methylation (**Fig EV5B**) (Roy *et al*, 2021; Wallner *et al*, 2016; Vento-Tormo *et al*, 2016). This effect was mirrored in DMR modeling, which identified 23,297 hypo and 3,660 hyper significant DMRs (Δ %5mC > ± 5%) among CS samples (**Fig 7E-F, Table EV7**). While many of these changes quite were small, a substantial number of sites (3,279 hypo and 1,163 hyper DMRs) exhibited a change DNA methylation greater than 10% 5mC, which is expected to greatly impact their regulatory activity.

Phylogenetic and principal components analyses revealed discrete clustering of DNA methylation patterns for each condition (**Fig 7D, Fig EV5C**). There was a strong overlap in DMRS between CS d6 and d7 (35% of hypo and 38% of hyper DMRs), although far more hypo DMRs were identified in CS d7 monocytes, corresponding with the greater degree of genome-wide hypomethylation observed in these cells (**Fig 7E**). Interestingly, nearly half of hyper DMRs were unique to CS d6 monocytes, suggesting the transient enrichment of a unique monocytic subtype cell in bone marrow during sepsis onset. Focusing on exhaustion DMRs, we noted significant DMRs at several regulatory sites previously implicated in our *ex vivo* model system, including enhancers for *Klf4*, *Nrp2*, and *Foxp1* (**Fig7F-G**). Hypomethylation was also observed at the *Socs3* distal enhancer and *Cfb* promoter in CS d6 and d7 cells, consistent with their sustained transcriptional activation at these time points (**Table EV7**). Furthermore, although the primary *Plac8* promoter exists in a lowly methylated state in undifferentiated bone marrow monocytes, we observed progressive hypomethylation at several of its intragenic regions in CS d6 and d7 cells, including an upstream alternative transcription start site. In total, of the 1,794 DMR-linked genes identified in our *ex vivo* BMMC sepsis model, roughly half were linked to significant DMRs *in vivo*, with 27% exhibiting strong (> 10% Δ5mC) differential methylation in CS monocytes (**Table EV1, Table EV7**). These results both support the involvement 5mC reprogramming in sepsis progression and validate our *ex vivo* model system as a tractable experimental paradigm for identifying key regulators of monocyte exhaustion.

Next, we annotated DMR probes in CS samples using sesame’s KYCG function (**Table EV8**). As with *ex vivo* LPS-treated BMMC samples, DMRs were heavily enriched at enhancer sites, a feature further corroborated by chromHMM analysis (**Fig EV5D**). Among CS d6 and d7 hyper DMRs, we noted strong enrichment for the tissue signature “monocyte hypomethylated”, again reflecting the disrupted methylome observed in BMMCs exhausted *ex vivo*. A survey of transcription factor binding motifs enriched at CS d6 and d7 DMRs identified IRF family members IRF1 and IRF8 among hyper DMRs, similar to what was observed *ex vivo*. By contrast, Wnt signaling factor MLLT3 was enriched among hypo DMRs in CS d6 and d7 monocytes, an inverse of the pattern observed in *ex vivo* cells; we also noted a strong enrichment of FOSL2 binding sites among hypo DMRs that was not evident *ex vivo* (**Fig EV5E, Table EV8**). Given how well DMRs were otherwise recapitulated *in vivo*, the implications of such altered transcription factor motif enrichment patterns remain unclear.

Gene ontology enrichment analysis of DMR-linked genes in CS d6 and d7 samples detected numerous regulatory pathways linked to sepsis progression, including “regulation of actin cytoskeleton”, “MAPK signaling”, “signaling by Rho GTPases”, “IL-3 signaling”, and “HIF1 signaling” among hypo DMRs and (**Table EV9**) (Weber *et al*, 2015). Interestingly, among hyper DMRs in CS d6 samples, we noted an enrichment for “phospholipase D signaling” and “phosphatidylinositol signaling”, pathways that have been recognized for their involvement in sepsis-associated lung injury and encephalopathy (Serantes *et al*, 2006; Recknagel *et al*, 2012; Lee *et al*, 2015; Abdulnour *et al*, 2018). We also noted an enrichment of “mitochondrial ABC transporters” among CS d7 hyper DMRs, consistent with the metabolic stress we and others have demonstrated in septic monocytes (Wasyluk & Zwolak, 2021).

Finally, to characterize the long-term changes in DNA methylation in bone marrow monocytes following sepsis resolution, we analyzed DNA methylation in purified bone marrow monocytes collected from mice after one week of sepsis recovery (CS d12). In contrast to the hypomethylation observed at CS d6 and d7, genome-wide DNA methylation in CS d12 monocytes normalized to levels observed in CTRL mice (**Fig EV5B**). Phylogenetic and principal components analyses of 5mC patterns in CS d12 monocytes likewise demonstrated a high degree of similarity with CTRL cells, indicating most of the changes observed during the acute sepsis had resolved during the recovery phase (**Fig 7D, Fig EV5C**). Despite this, we still observed 97 hypo and 203 hyper DMRs relative to CTRL (**Fig 7E**). Whereas most hypo DMRs were shared with sites observed in CS d6 and d7 monocytes (81%), more than half of hyper DMRs were unique to CS d12 cells. Again, sesame KYCG annotation and chromHMM analysis revealed an enrichment of CS d12 DMRs at enhancers (**Fig EV5D, Table EV8**). Among these sites were enhancers linked to critical immune genes such as *Il10*, *Socs3*, and *Nrp2* (**Fig 7G**). GO analysis detected an enrichment for numerous immune pathways among CS d12 DMR-linked genes, including “regulation of actin cytoskeleton”, “phospholipase D signaling”, “signaling by Rho GTPases”, “MAPK signaling”, and “IL-3 signaling” (**Table EV9**). Strikingly, there was profound enrichment for the FOSL2 binding motif among hypo DMRs (30% of hypo DMRs), implicating the transcription factor as an important regulator of long-term epigenetic changes in immune challenged monocytes (**Fig EV5C**). Taken together, these results demonstrate that while 5mC reprogramming seems to play a greater role in regulating monocyte behavior during acute sepsis, changes in DNA methylation contribute to long-term innate immune memory in monocytes well after sepsis resolution.

## DISCUSSION

Innate immune memory (IM) is essential for leukocyte homeostasis, facilitating long-term transcriptional reprogramming in response to immune challenge in a pathogen-and concentration-specific manner (Ifrim *et al*, 2014; Netea *et al*, 2011; Quintin *et al*, 2014; Netea *et al*, 2020a). These altered memory states can have far reaching effects on innate immune activity. In sepsis, IM transcriptional reprogramming results in disrupted myeloid cell behavior as late as three months after sepsis recovery, challenging traditional notion of adaptive immunity as the *de facto* driver of long-term immune memory (Shalova *et al*, 2015; Bomans *et al*, 2018). IM has also been implicated in long-term sepsis-induced immunoparalysis, complications from which facilitate persistent and nosocomial infection leading to death in one-third of sepsis patients (Hamers *et al*, 2014; Baudesson de Chanville *et al*, 2020; Gentile *et al*, 2012; Prescott & Angus, 2018; Netea *et al*, 2016).

Central to IM are alteration to the epigenome, most notably in the form of histone modifications. Major changes to active histone marks H3K27ac and H3K4me1/3 have been demonstrated under both IM tolerance and training in a wide variety of biological contexts, including sepsis, endotoxin tolerance, Bacille Calmette-Guérin (BCG) vaccination, wound repair, and systemic sclerosis (Davis *et al*, 2019; Saeed *et al*, 2014; Kleinnijenhuis *et al*, 2012; Foster *et al*, 2007; Van Der Kroef *et al*, 2019; Novakovic *et al*, 2016). Importantly, these changes are not merely correlative, as pharmacological inhibition of H3K4 and H3K27 histone readers and writers have been clearly demonstrated to block IM transcriptional reprogramming (Novakovic et al, 2016; Nicodeme et al, 2010; Bekkering et al, 2014; Ifrim et al, 2014; Van Der Kroef et al, 2019).

Less well understood is the role epigenetic regulation in the form of DNA methylation plays in IM. Focused studies on the *Tlr4* promoter in CD14^+^ human monocytes demonstrated RFX1-mediated recruitment of DNMT1 to drive local DNA methylation and gene repression, a process that is impaired in low-density lipoprotein exposure in coronary artery disease patients (Du *et al*, 2019). At the genomic level, following BCG vaccination, stable differences in DNA methylation patterns were observed in peripheral blood mononuclear cells (PBMCs) of responders versus non-responders up to 8 months post-vaccination; similar differences were also observed in adaptive natural killer cell subsets in response to CMV infection (Verma *et al*, 2017; Schlums *et al*, 2015). By contrast, Novakovic et al. 2016 found no evidence for the involvement of DNA methylation in β-glucan-driven trained immunity, although significant differences were observable in LPS tolerized macrophages (Novakovic *et al*, 2016) Similarly, multiomics integration identified a link between DNA methylation and endotoxin tolerance via a transcriptional signature attuned to cholesterol biosynthesis during the acute stage of pneumonia (Brands *et al*, 2021). Perhaps most compellingly, PBMCs collected from septic patients exhibited altered DNA methylation profiles that correlated with inflammatory cytokine secretion and organ dysfunction (Lorente-Sorolla *et al*, 2019). In a follow-up study, the authors found evidence for a JAK/STAT signaling axis underlying broad DNA demethylation in LPS tolerized monocytes (Morante-Palacios *et al*, 2021). However, whether DNA methylation reprogramming contributes to pathogenic features of monocyte exhaustion during sepsis, as well as information on the long-term implications for sepsis-driven 5mC alterations, remained unknown.

In our present study, we utilized a recently validated *ex vivo* mouse sepsis model to more fully characterize the involvement of DNA methylation in IM and sepsis pathogenesis (Pradhan *et al*, 2021). In addition to affording researchers an experimentally tractable method for conducting mechanistic and therapeutic studies in primary mouse monocytes, this model system also recapitulates the paradoxical pro-inflammatory, anti-immune behavior characteristic of exhausted monocytes during acute sepsis. Our results clearly demonstrate that changes to the 5mC landscape of exhausted monocytes underlies much of this pathogenic behavior. Differential methylation at gene regulatory sites for transcription factors (e.g. *Foxp1*, *Cebpa/g*, *Runx3*) and immune effectors (e.g. *Socs3*, *Trem2*, *Cfb*) correlated strongly with their altered transcriptional state in exhausted monocytes, and pharmacological inhibition of DNMT activity restored immune function at the levels of transcriptional and cell surface marker expression. Most of these changes occurred at gene enhancers and flanking transcription start sites, correlating strongly with altered H3K27ac peaks in these regions previously observed in an exhausted monocyte dataset (Naler *et al*, 2022). While the relationship between altered DNA methylation patterns and H3K4 methylation was not also explored in this present study, previous research has shown that H3K4me4 directly antagonizes the recruitment of DNMT3A and its cofactor DNMT3L, thereby inhibiting *de novo* DNA methylation (Ferry *et al*, 2017; Otani *et al*, 2009; Ooi *et al*, 2007). Given the extent of genome-wide H3K4me1/3 reprogramming normally observed during sepsis and LPS tolerization, it is reasonable to speculate that DNA methylation serves to reinforce changes to the H3K4 methyl landscape in exhausted monocytes (Saeed et al, 2014; Foster et al, 2007; Novakovic et al, 2016).

Perhaps one of the most interesting hits among DMRs in exhausted monocytes was strong hypomethylation in the proximal promoter region of *Plac8*. Although *Plac8* was first identified as a leukocyte inhibitory factor enriched in mouse uterus and placenta, recent studies have demonstrated an important role for *Plac8* during severe immune challenge (Galaviz-Hernandez *et al*, 2003). In fact, *Plac8* was identified as the top enriched gene in pathogenic monocyte subtypes based on single-cell analyses of sepsis and Covid-19 patients (Schulte-Schrepping *et al*, 2020; Reyes *et al*, 2020). Furthermore, two recent studies demonstrated *Plac8* expression as essential for coronavirus infection by regulating either viral entry or transcription in host cells (Tse *et al*, 2022; Ugalde *et al*, 2022). Whereas previous studies have implicated *Plac8* hypomethylation in endothelial colony-forming cells as a driver for *Plac8* upregulation and downstream transcriptional alterations in gestational diabetes mellitus, to our knowledge our study is the first to report DNA methylation as potential driver for *Plac8* monocytic dysregulation during sepsis progression (Blue *et al*, 2015). Given the dual roles of PLAC8 performs as both a regulator of autophagosomal-lysosomal fusion and a bona fide transcription factor through interactions with C/EBPβ, the mechanism by which PLAC8 impacts immune behavior in exhausted monocytes remains an active area of investigation (Kinsey *et al*, 2014; Jimenez-Preitner *et al*, 2011).

Single-cell modeling of major regulons during monocyte exhaustion identified several transcription factors previously implicated in sepsis progression, including HIF1A, CEBPB, CEBPD, and NF-KB2 (Reyes *et al*, 2020; Schulte-Schrepping *et al*, 2020; Avendanõ-Ortiz *et al*, 2018b; Kang *et al*, 2016). Furthermore, motif analysis for transcription factor binding sites demonstrated an enrichment for many of these factors at DMRs, suggesting 5mC reprogramming in exhausted monocytes helps to direct and stabilize their regulatory activity. One of the more interesting observations from these analyses was evidence for Wnt signaling inhibition during monocyte exhaustion, supported by hypermethylation of MLLT3 binding sites in LPS-treated cells and TCF7L2 signaling activity specific to non-classical PBS control monocytes. Up to this point, the role of Wnt signaling in monocytes has been poorly characterized. Previous studies demonstrated that Wnt signaling promotes monocyte transendothelial migration and cellular adhesion (Tickenbrock *et al*, 2006; Lee *et al*, 2006). It has also been shown that Wnt signaling inhibits PU.1-mediated gene regulation to block monocyte-macrophage differentiation, although that effect was not recapitulated in our study likely due to the low concentration of Wnt Agonist 1 tested (Sheng *et al*, 2016). Instead, activation of Wnt signaling during repetitive LPS stimulation largely antagonized the exhaustion phenotype, redistributing monocyte subtypes from the pro-inflammatory Ly6C^high^ to the pro-resolving Ly6C^low^ fate and downregulating the expression of major exhaustion genes such as *Plac8*, *Hif1a*, *Runx3*, and *Cd38*. Furthermore, Wnt signaling activation restored DNA methylation to control levels at several key regulatory features, including the *Plac8* promoter, *Cebpa/g* enhancer, and *Klf4* enhancer. Taken together, these results support Wnt signaling as a major regulatory node in monocytes during pathogenic inflammation and prospective target for therapeutic intervention.

Several previous studies have demonstrated the translational potential for innate immune training as both a prophylactic measure to mitigate infection risk and regenerative feature to treat immunoparalysis (van der Meer *et al*, 2015; Netea *et al*, 2016; Bekkering *et al*, 2015; Netea *et al*, 2020b, 2020a). Chief among these therapeutic agents are fungal derivative β-glucan and BCG derivatives such as TDM, which have been shown to limit disease risk (most notably in SARS-CoV-2 infection) and reduce the severity of symptoms after infection occurs (Quintin *et al*, 2012; Rodriguez *et al*, 2019; Netea *et al*, 2020b; Freyne *et al*, 2015; Moorlag *et al*, 2019; Escobar *et al*, 2020; Netea *et al*, 2021; Rivas *et al*, 2021; Kong *et al*, 2021). Furthermore, treatment of LPS- tolerized macrophages with β-glucan restores H3K27ac levels at tolerized genes, supporting the use of innate immune training as a means to modulate those epigenetic changes underlying IM (Novakovic *et al*, 2016). Given that the influence of immune training on DNA methylation in LPS-stimulated cells has been previously untested, we cultured exhausted monocytes in the presence of BCG-derivative TDM to measure its impact on 5mC reprogramming (Madonna *et al*, 1989b, 1989c). Strikingly, TDM intervention had a profound effect on DNA methylation patterns in exhausted monocytes, greatly limiting the number of hypermethylated regions observed in these cells and restoring 5mC levels at regulatory sites for critical immune modulators such as *Pim1*, *Foxp1*, *Socs3*, *Nfatc2*, and *Cd83*. Thus, immune training with BCG-derivatives has the potential to promote healthy epigenetic profiles at the level of both histone modifications and DNA methylation, further supporting the therapeutic potential of these agents.

In recent years, the importance of epigenetic alterations in long-term IM have been a subject of intense debate. In support for an epigenetic etiology, disrupted H3K4me3 levels at the *Il1b*, *Il12*, and *Il23* gene promoters were reported in murine bone marrow stem and progenitor cells 4 weeks after sepsis recovery (Davis *et al*, 2019). In a separate study, increased levels of H3K4me3 were visible at the *Tnfa*, *Il6*, and *Tlr4* promoters four months after BCG vaccination in human patients (Kleinnijenhuis *et al*, 2012). Similar effects have also been observed in tissue resident macrophages (Yao *et al*, 2018; Roquilly *et al*, 2020). By contrast, a recent time course analysis of H3K27ac and H3K4me1/3 in immortalized bone marrow-derived macrophages following β-glucan exposure found that many of these changes are transient, with most differential peaks resolving within two weeks of stimulation (Sun *et al*, 2023). Instead, the authors advocate for sustained transcription factor activity as essential for the transmission of stimulus-induced epigenetic changes across cellular divisions. In our own study, we found evidence for both models in DNA methylation reprogramming of monocytes during sepsis progression. As with histone modifications, the altered 5mC profile observed at early time points following cecal slurry injection were largely resolved one week after sepsis recovery, indicating DNA methylation alteration may play a more important role in regulating monocyte behavior during the acute phase of sepsis. However, stable changes in DNA methylation were noted at several hundred immune loci, with differential methylation at some sites (e.g. *Nrp2* enhancer, *Fgfbp3* promoter) actually intensifying during this period. That many of these changes were also observed in our *ex vivo* exhaustion model argues for a cell autonomous role in DMR establishment; as to whether paracrine signaling cues are needed to potentiate these changes will require further investigation.

In sum, our findings conclusively support a role for DNA methylation in IM and monocyte exhaustion under septic conditions. We also note the translational potential for pharmacological intervention in the form of Wnt agonists, DNA demethylating drugs, and BCG-derived immune training agents in the resolution of aberrant gene expression and 5mC patterning in exhausted monocytes. Given the strong association between DNA methylation and pathogenic drivers such as *Plac8*, DNA methylation merits equal consideration to histone modifications for the role it plays in shaping the epigenetic landscape of IM.

## MATERIALS & METHODS

### Animal husbandry

All experiments were approved by the Institutional Animal Care and Use Committee (IACUC) of Virginia Tech in accordance with the U.S. National Institutes of Health Guide for the Care and Use of Laboratory Animals. Wild-type (WT) C57BL/6 mice (originally from The Jackson Laboratory), and *Ticam2^-/-^* mice (provided by Dr. Holger Eltzschig at University of Texas Houston) are regularly bred and maintained in our laboratory. Mice were housed in a pathogen-free facility with 12-12 hour light-dark cycles and free access to water and standard chow.

### In vitro cell culture

*In vitro* bone marrow monocyte (BMMC) LPS exhaustion experiments were performed as previously described (Pradhan *et al*, 2021). Briefly, bone marrow cells were harvested from 6- to 8-week-old C57BL/6 WT or *Ticam2^-/-^* female mice, seeded at a density of 3×10^5^ cells/cm^2^, and cultured in complete RPMI 1640 media (10% fetal bovine serum, 1% penicillin-streptomycin, 1% L-glutamine) supplemented with 10 ng/mL M-CSF (PeproTech). Cells were cultured for 5 days at 37°C in a humidified 5% CO_2_ atmosphere under continuous high-dose 100 ng/mL lipopolysaccharide (LPS; Sigma-Aldrich) stimulation or PBS control conditions, with fresh media changes at days 2 and 4 of culture (Geng *et al*, 2016). For Monensin A (MonA; Sigma Aldrich) and Wnt agonist 1 (Cayman Chemical) experiments, cells were treated with low-dose MonA (50 nM) or Wnt agonist 1 (1 or 5 µM) on days 2 and 4 of cell culture. For 5-azacytidine (5-aza; Stem Cell Technologies) and trehalose 6,6’-dimycolate (TDM; invivoGen) experiments, cells were maintained in 5-aza (50 or 250 nM) or TDM (10 µg/mL) for the full 5 day LPS exhaustion time course.

### Cecal slurry preparation and injections

Sepsis was induced by intraperitoneal injection of cecal slurry in 6- to 8-week-old C57BL/6 WT male mice as previously described (Starr *et al*, 2014). Briefly, whole ceca were dissected from 12-week-old C57BL/6 mice euthanized by cervical dislocation. Cecal contents were extracted and mixed with sterile water (250 µg/µL), sequentially filtered through 860 µm and 190 µm mesh strainers, mixed with an equal volume of 30% glycerol in PBS (final concentration 125 µg/µL), and placed in −80°C storage. For cecal slurry injections, frozen cecal slurry (CS) stock was rapidly thawed in 37°C water bath and injected (0.9 mg/g mouse weight) into the peritoneal cavity of recipient mice. After 5 days, mice were injected with a second 0.9 mg/g dose of CS, and then euthanized by cervical dislocation 1, 2, or 7 days later. Bone marrow cells were collected as previously described and used for flow cytometry analysis or LS column purification (Miltenyi) of bone marrow monocytes following the manufacturer’s protocol.

### Flow cytometry and fluorescence activated cell sorting (FACS)

Mouse bone marrow cells or day 5 cultured BMMCs were washed with PBS, harvested, and blocked in 1:100 Fc block (BD Biosciences) for 20 minutes at 4°C. Cells were then incubated for 30 minutes at 4°C with one of three separate antibody panels. For panel 1, cells were stained with fluorochrome-conjugated antibodies against Ly6C (PE-Cy7; Biolegend #128017), CD11b (APC-Cy7; Biolegend #101226), CD86 (FITC; Biolegend #105006), PD-L1 (APC; Biolegend #124312), and CD38 (PE; Biolegend #102708). For Panel 2, cells were stained with antibodies against Ly6C (Pacific Blue; Biolegend #128014), CD11b (BV650; BD Biosciences #563402), Ly6g (PE-Cy5.5; Elabscience #E- AB-F1108I), CXCR2 (Alexa Fluor 488; R&D Systems # FAB2164G), CD68 (APC-Fire750; Biolegend #137041), CD172a (BV510; BD Biosciences #740159), CX3CR1 (PE-Cy7; Biolegend #149016), F4/80 (BV711; BD Biosciences #565612), CD38 (BV750; BD Biosciences #747103), PD-L1 (BV421; BD Biosciences #564716), MARCO (Alexa Fluor 647; R&D Systems #FAB29561R), FAS (BV605; Biolegend #152612), MHCII (PerCP; Biolegend #107624), CD168 (PE; Novus #NBP1-76538PE), and TREM2 (Alexa Fluor 700; R&D Systems #FAB17291N). For panel 3, cells were stained with only Ly6C (PE-Cy7; Biolegend #128017) and CD11b (APC-Cy7; Biolegend #101226). Cells were then washed with FACS buffer (HBSS with 2% fetal bovine serum) and resuspended in FACS buffer containing propidium iodide (PI; 1:2000; Invitrogen). Panel 1 samples were measured with a FACS Canto II (BD Biosciences), panel 2 samples were measured with an Aurora 4L 16V-14B-10YG-8R (Cytek), and panel 3 samples were sorted by FACS using an SH800S Cell Sorter (Sony).

### NAD^+^ assay

An Amplite Fluorimetric cADPR-Ribose Assay Kit (AAT Bioquest) was used to determine NAD^+^ in cultured BMMCs according to the manufacturer’s protocol. NAD^+^ levels were quantified using a Cytation 3 Cell Imaging Multi-Mode Reader (BioTek).

### Quantitative real-time PCR (qRT-PCR)

Total RNA was extracted using an RNeasy Plus Mini Kit (QIAGEN) and reverse transcribed using a High-Capacity cDNA Reverse Transcription Kit (Applied Biosystems) according to the manufacturer’s protocol. qRT-PCR was performed using Power SYBR Green Master Mix (Thermo Fisher) on a CFX96 Real-Time System C1000 Thermal Cycler (Bio-Rad). Relative expression levels were determined using the Pffafl method normalized to the geometric mean of genes *Ube2l3*, *Oaz1*, and *Nktr*, selected based on scRNA-seq data demonstrating their high, equivalent expression levels across all cell types tested (Pradhan *et al*, 2021)

### Infinium Mouse Methylation BeadChip Array

Following 5 days of LPS or PBS control stimulation, CD11b^+^ BMMCs were FACS sorted into Ly6C-low, -intermediate, and -high pools. Genomic DNA from each pool was prepared using a DNeasy Blood & Tissue Kit (QIAGEN) and bisulfite (BS)-treated using an EpiTect Bisulfite Kit (QIAGEN). For Infinium Mouse Methylation BeadChip (Illumina) arrays, 500 ng of BS-treated DNA was processed and hybridized to individual array wells according to the manufacturer’s protocol. BeadChip array signal was measured using the iScan System (Illumina). All samples were processed and run at the Children’s Hospital of Philadelphia Center for Applied Genomics.

### Bisulfite next-generation sequencing

A modified BisPCR^2^ workflow was used to validate differentially methylated regions of interest identified on the Infinium array (Bernstein *et al*, 2015). Genomic DNA was subjected to bisulfite conversion as described above and then used as a template for target enrichment using a PyroMark PCR Kit (QIAGEN). Amplified regions were pooled for column purification (4-7 regions per pool, for a total of 150ng), and the purified pools were barcoded with Illumina indexing primers using a Multiplex PCR Kit (QIAGEN). All indexed pools for each sample were then pooled during column purification, after which library quality was determined using Tapestation DNA ScreenTape analysis (Agilent) at the Fralin Life Sciences Institute at Virginia Tech Genomics Sequencing Center. Finally, all indexed libraries were pooled with 6% PhiX spike-in DNA and sequenced using a MiSeq Reagent Nano Kit v2 (500 cycles) (Illumina) at the Fralin Genomics Sequencing Center.

### Quantification and Statistical Analysis

#### Statistics

General statistical analyses were performed using Prism 9 (Graphpad). Comparisons between two groups were performed using a two-tailed t-test based on the normal distribution of the data. Comparisons between three or more groups were performed using one-way ANOVA adjusted for Tukey’s multiple comparisons, while paired comparisons of PBS control and LPS exhaustion conditions were corrected using Šidák based on the assumption that each comparison is independent of the other. Pearson correlation analyses for DNA methylation against gene expression or H3K27ac signal, as well as Mood’s nonparametric median test for differences in genomic 5mC distributions between cecal slurry samples, were performed in R (R Core Team, 2021). Sample sizes were chosen based on variation observed in previous BMMC and CS datasets from our laboratory (Lin et al, 2020; Pradhan et al, 2021; Lee et al, 2021; Geng et al, 2021; Naler et al, 2022).

#### Infinium BeadChip Array analysis

Raw Infinium IDAT files were processed into corrected beta values using the openSesame pipeline (1.14.2; default parameters) (Zhou *et al*, 2018). Differentially methylated regions (DMRs) between PBS- and LPS-treated BMMCs were identified using the sesame DML function, with Ly6C cell surface expression (low, intermediate, high) being used as a covariate. Region annotations for DMRs were obtained using the annotatePeaks function of the Rpackage ChIPSeeker (1.22.1; parameter: TxDb=TxDb.Mmusculus.UCSC.mm10.knownGene) (Yu *et al*, 2015). Additional Infinium BeadChip probe annotations, including transcription factor motif enrichment, chromatin state discovery and characterization (ChromHMM), and tissue signatures, were obtained using the sesame KYCG function (Zhou *et al*, 2018). For HOMER motif enrichment, differentially methylated probes sites were expanded to non-overlapping 500 bp windows (+/- 250 bp from probe CpG) and analyzed using the findMotifs.pl program relative to the Infinium Methylation BeadChip array background (Heinz *et al*, 2010). For gene ontology enrichment analysis, differentially methylated probes were matched to their nearest gene using sesameData_getGenesByProbes, and ontology enrichment was performed using the g:Profiler2 package (0.2.1) (Peterson *et al*, 2020). Average DNA methylation signal and unsupervised clustering heatmaps were prepared using the Rpackage pheatmap (1.0.12; https://www.rdocumentation.org/packages/pheatmap; clustering_method = “average”), and average signal violin plots were prepared using the Rpackage vioplot (0.4.0; https://github.com/TomKellyGenetics/vioplot). Principal components analysis (PCA) and Moods nonparametric median tests were performed using the R stats package.

#### Chromatin immunoprecipitation (ChIP)-seq analysis

BMMC H3K27ac ChIP-seq data (GEO accession: GSE168190) quality control, normalization, and peak calling was performed as previously described (Data ref: Naler et al, 2022). H3K27ac peaks overlapping DMRs in LPS-treated BMMCs were identified using BEDTools (2.30.0; function: intersect) (Quinlan & Hall, 2010). Heatmaps and metaplots for H3K27ac signal at DMRs were generated using deepTools (3.5.2), and correlation plots for differential H3K27ac peaks at DMRs were generated using ggplot2 (3.4.0) (Ramírez et al, 2016; Ginestet, 2011).

#### Bisulfite next-generation sequencing analysis

Sequenced reads were trimmed using Trim Galore (0.6.7; http://www.bioinformatics.babraham.ac.uk/projects/trim_galore) and mapped with Bismark (0.22.3; https://www.bioinformatics.babraham.ac.uk/projects/bismark) in paired-end mode. Non-deaminated reads were filtered out based on the presence of ≥ 3 consecutive instances of non-CG methylation (function: filter_non_conversion ; parameters: --paired --consecutive). Bedgraph files were prepared using the Bismark Methylation Extractor to calculate percent methylation at each CpG with ≥ 100x coverage. Aligned reads were converted to Bigwig tracks for UCSC browser visualization (http://genome.ucsc.edu) (Kent et al, 2010).

#### Single-cell sequencing analysis

Single-cell sequencing data from WT and *Ticam2^-/-^* BMMCs (Genebank accession: GSE182355) were processed using 10x Genomics Cell Ranger (3.1.0), with reads mapped using the count pipeline and pre-built reference genome refdata-gex-mm10-2020-A and GTF from GENCODE vM23 (GRCm38) (Data ref: Pradhan et al, 2021). Downstream analyses were performed in R using Seurat (4.3.0) (Stuart *et al*, 2019). Low quality cells were filtered out based on the following criteria: > 20% mitochondrial reads, < 2.5% ribosomal reads, or < 200 unique genes for a given cell. Following the removal of doublets and a small population of *Igcl1/2*-positive B cells, 2483 high-quality BMMC cells remained for downstream analysis. For genes expressed in > 2 cells, expression was normalized (normalization.method = “LogNormalize”, scale.factor = 10,000), and principal component analysis (PCA) was performed using the 2000 most variable genes (selection method = “vst”). The jack-straw method (num.replicate = 100, dims = 50) was used to identify suitable dimensionality for cell clustering (42 dimensions; resolution = 0.6). Cell clusters were visualized with UMAP and annotated based on marker gene expression (FindMarkeres; min.pct = 0.25, logfc.threshold = 0.25).

#### SCENIC analysis

To identify BMMC single-cell cluster transcription factor regulatory modules, the SCENIC workflow was performed as previously described (Aibar et al, 2017). Expression matrices were prepared for filtered genes expressed in at least 1% of cells and with > 200 counts. SCENIC-derived AUC values for filtered transcription factor modules were visualized via UMAP in Seurat (Stuart *et al*, 2019). Regulon specificity scores (RSS) for each transcription factor module were plotted for all BMMC cell clusters (calcRSS; thr = 0.01, zThreshold = 2).

#### Flow cytometry analysis

All flow cytometry data were analyzed with FlowJo (BD Life Sciences). For *ex vivo* BMMC experiments, mean fluorescence intensity (MFI) values for each antibody were normalized to the average MFI value for WT PBS control cells on a given experimental day. For FlowSOM analysis of bone marrow cells, individual CTRL and CS replicates were downsampled to 15,000 (CTRL) or 10,000 (CS) cells using the DownSampleV3 plugin, for a total of 120,000 cells per condition. Next, cells were clustered using the FlowSOM plugin (15 Metaclusters; Set Seed = 3). All clusters with < 0.5% cells represented were removed from the analysis (Pop0, Pop4, and Pop10), as well as one cluster with poor mapping consistency (Pop13). Cell clusters were visualized using the UMAP plugin.

## ACKNOWLEDGMENTS

The authors would like to thank Yao Zhang for assistance with mouse dissections and Feng Xu for mouse colony management. We would also like to thank technicians at the Fralin Life Sciences Institute at Virginia Tech Genomics Sequencing Center and Children’s Hospital of Philadelphia Center for Applied Genomics for their service and guidance in DNA methylation array- and sequencing-based approaches. This work was supported by the National Institutes of Health: 5TL1DK132771 (B.A.C.) and R01-AI-172133 (L.L.). Additional funding support was provided by the Virginia Presidential Postdoctoral Fellowship Program and its sponsors, the Office of Research and Innovation, Fralin Life Sciences Institute, and Institute for Critical Technology and Applied Science at Virginia Tech (B.A.C.).

## AUTHOR CONTRIBUTIONS

**Blake A. Caldwell:** Conceptualization; data curation; formal analysis; investigation; visualization; methodology; validation; software; funding acquisition; writing – original draft; writing – review and editing. **Yajun Wu:** Conceptualization; investigation; formal analysis; methodology; writing – review and editing. **Jing Wang:** Investigation; methodology; writing – review and editing. **Liwu Li:** Conceptualization; resources; project administration; supervision; funding acquisition; writing – review and editing.

## THE PAPER EXPLAINED

### Problem

Monocyte dysregulation is thought to underlie the most pathogenic features of sepsis and Covid-19 infection, notably as a driver for the “cytokine storm” phenomenon. However, the molecular mechanism for this dysregulation, in particular how it contributes to long-term immune complications in survivors, remains poorly defined.

### Results

Here, we tested the contribution of epigenetic changes in the form of DNA methylation to acute and long-term monocyte dysregulation during sepsis progression. Altered DNA methylation was present at regulatory features for numerous critical immune genes (e.g. *Plac8*, *Socs3*, *Cfb*), and resolution of these epigenetic patterns was associated with improved immune activity in septic monocytes.

### Impact

In addition to histone modifications, DNA methylation should be considered as a major epigenetic regulator of monocyte behavior and innate immune memory dynamics during septic challenge. Pharmacological intervention in the form of DNA methyltransferase inhibitors, Wnt signaling agonists, and innate immune training agents all represent viable therapeutic strategies to promote healthy monocyte behavior during severe immune challenge.

## DATA AVAILABILITY

The datasets produced in this study are available in the following databases:

- DNA methylation array data: Gene Expression Omnibus (pending)

## EXPANDED VIEW FIGURE LEGENDS

**Figure EV1.**
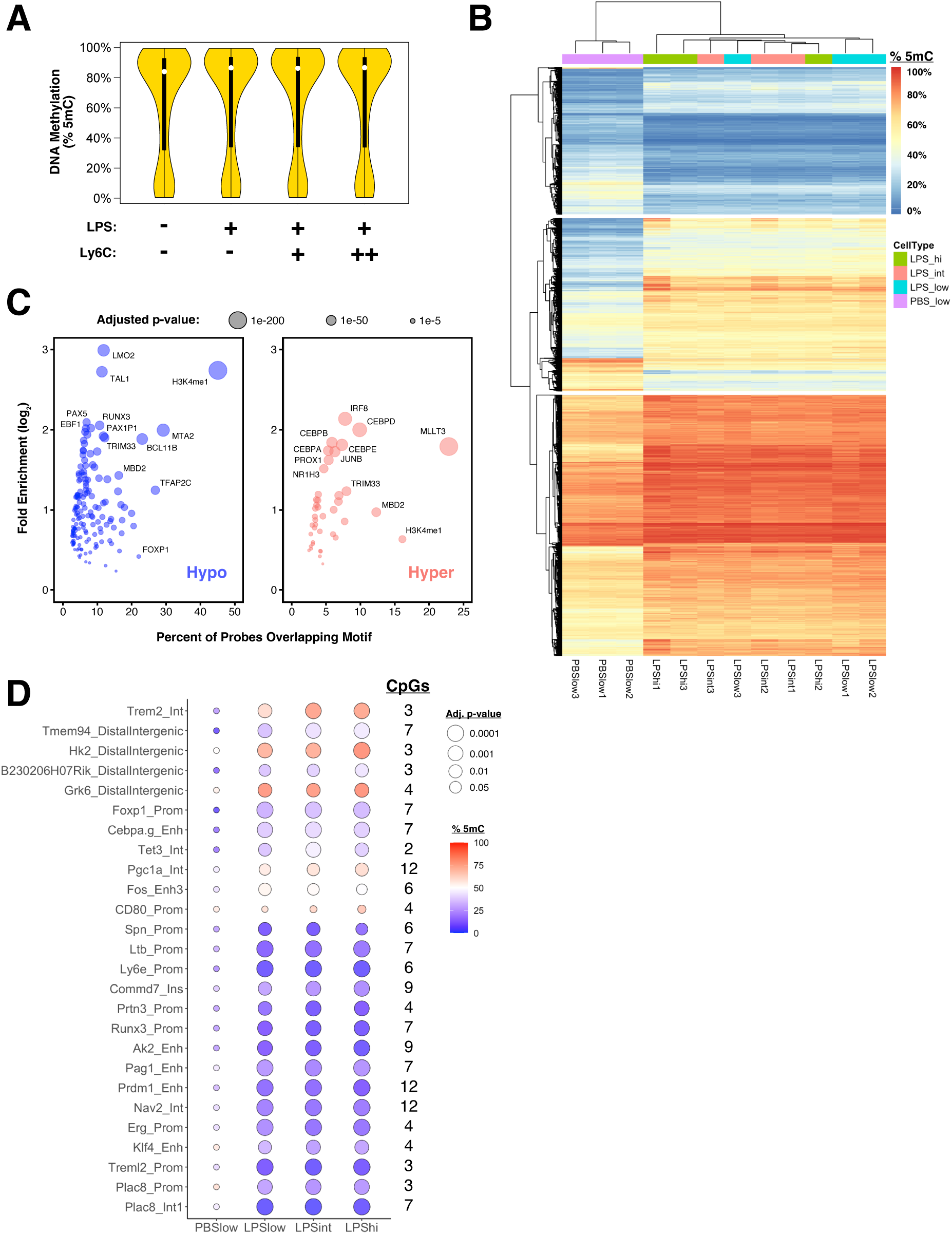
Extended analysis of DNA methylation in exhausted monocytes. **(A)** Violin plots of average DNA methylation levels at each probe CpG for the BMMCs under PBS control or repetitive-LPS stimulation, with box plots indicating medians, interquartile ranges, and 95% confidence intervals (n = 3 for each condition). **(B)** Correlation heatmap of DNA methylation at differentially methylated CpG probes following 5 days of PBS control or repetitive LPS stimulation (≥ 5% difference in % 5mC vs. PBS control; FDR < 10%; n = 3 for each condition). Columns represent individual biological replicates, and rows represent individual probe CpG sites. Clustering of replicates and probes was performed using the “Average” method, with dendrogram heights proportional to cluster distances. **(C)** Sesame Know Your CpG (KYCG) transcription factor (TF) binding motif analysis for hyper and hypo DMRs. **(D)** Heatmap of mean DNA methylation levels at MiSeq-validated DMRs (n = 3 for each condition; one-way ANOVA with Tukey multiple comparisons). Significant differences (p-adj. < 0.05) were observed at each DMR for all LPS monocyte subtypes with the exception of CD80 promtoer, *Fos* enhancer 3 (adj. p-value Ly6c^int^ and Ly6C^high^ = 0.06 and 0.13, respectively) and *Pgc1α* intron (adj. p-value Ly6C^low^ = 0.07).

**Figure EV2.**
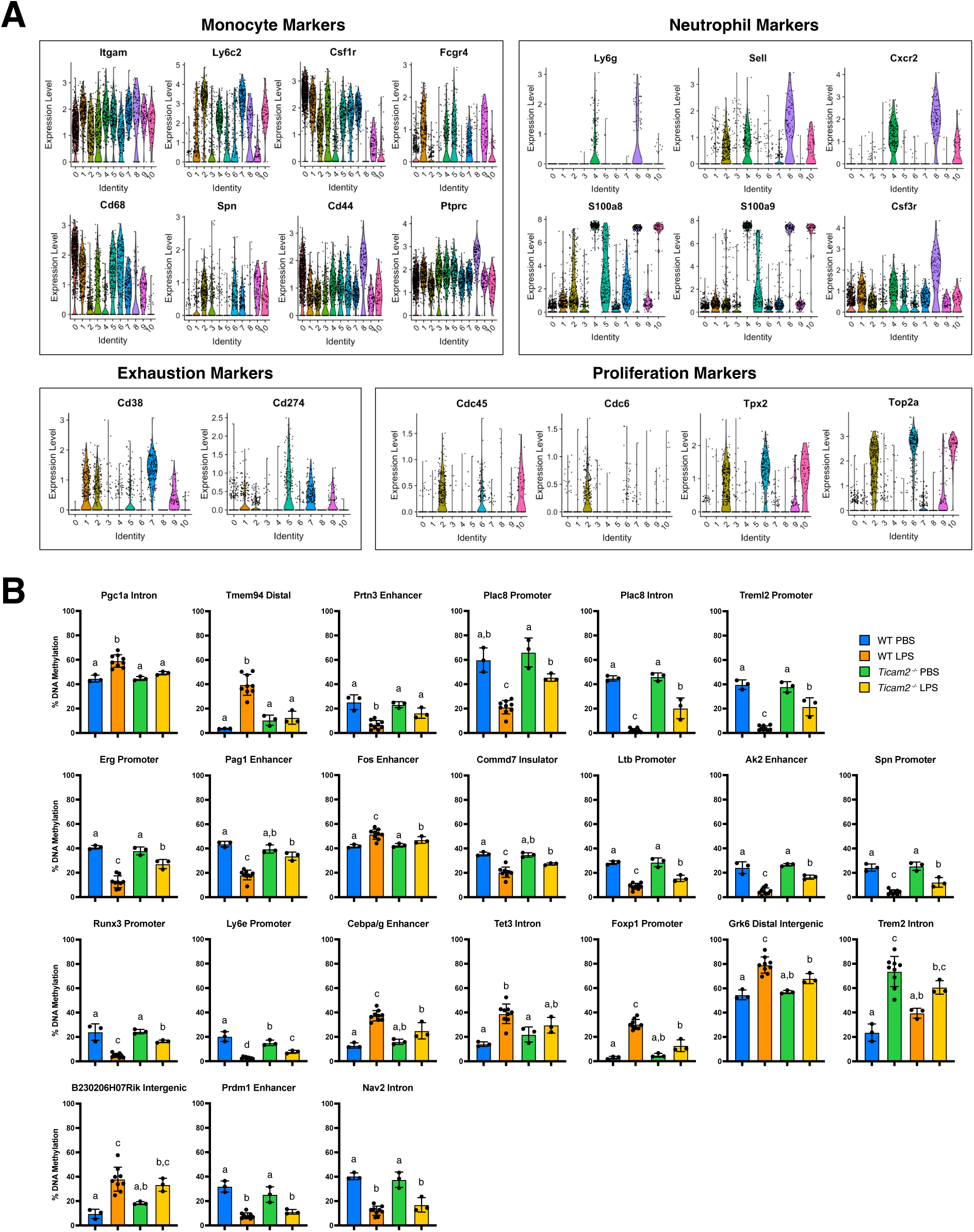
scRNA-seq marker gene expression and extended analysis of DNA methylation in exhausted monocytes. **(A)** Gene expression levels across cell clusters for various markers used to classify monocyte subtypes and phenotypes. **(B)** MiSeq DNA methylation levels at immune-relevant DMRs in WT and *Ticam2^-/-^*BMMCs following 5 days of PBS control or repetitive LPS stimulation. Bar graphs represent means ± STDs, with letters designating statistically distinct groups (1-way ANOVA with Tukey multiple comparisons; n=3-9). Full rescue DMRs are designated based on *Ticam2^-/-^*LPS DNA methylation levels being statistically different from WT LPS but not WT PBS, while partial rescue DMRs exhibit intermediate *Ticam2^-/-^*LPS DNA methylation levels statistically distinct from both WT PBS and WT LPS.

**Figure EV3.**
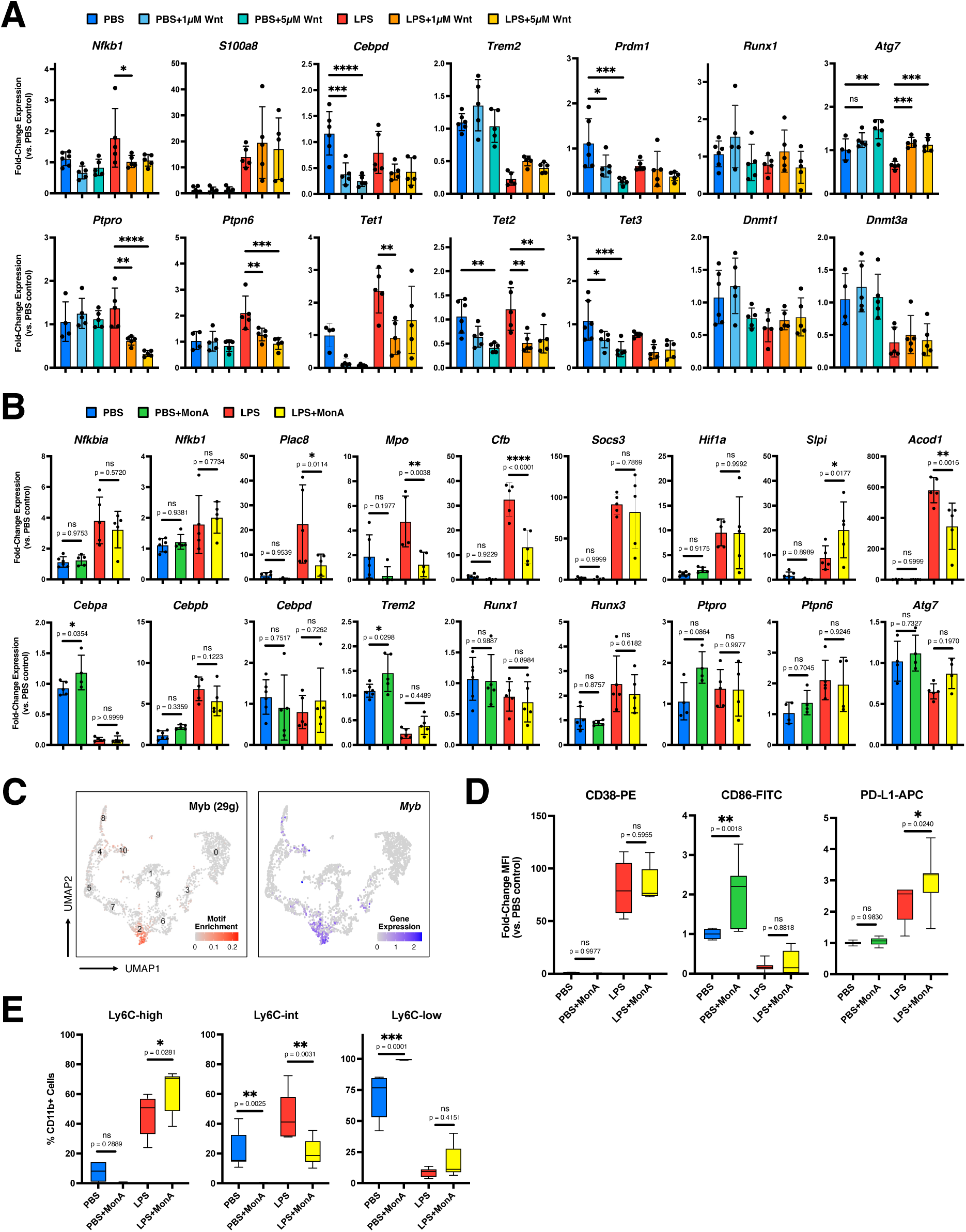
Extended analysis of Wnt signaling effect on monocyte exhaustion. **(A)** qRT-PCR for additional exhaustion markers in BMMCs under PBS control or repetitive LPS stimulation in the presence of different concentrations of Wnt Agonist 1. Expression levels were normalized to the geometric mean of *Ube2l3*, *Oaz1*, and *Nktr*. Data points represent independent experiments, with mean expression +/- STD indicated (n = 3-6; one-way ANOVA with Sidak’s multiple comparisons test; **** p-adj. < 0.0001; *** < 0.001; ** < 0.01; * < 0.05; ns not significant; differences are not significant unless otherwise specified; see Appendix for exact p-values). **(B)** qRT-PCR for exhaustion markers in BMMCs under PBS control or repetitive LPS stimulation in the presence of different concentrations of Wnt inhibitor Monensin A (MonA). **(C)** scRNA-seq UMAP feature plots for transcription factor Myb motif enrichment and gene expression in WT and *Ticam2^-/-^* BMMCs following PBS control or repetitive LPS stimulation. Numbers indicate different cell clusters, as outlined in Figure 2A. **(D)** Flow cytometry mean fluorescence intensity (MFI) for exhaustion markers relative to PBS control. Box plots indicate median MFI values (n = 7; one-way ANOVA with Sidak’s multiple comparisons test). **(E)** Population statistics for non-classical (Ly6C-low), intermediate (Ly6c-int), and classical (Ly6C-high) monocyte subtypes in cultured BMMCs (n = 7 ; one-way ANOVA with Sidak’s multiple comparisons test).

**Figure EV4.**
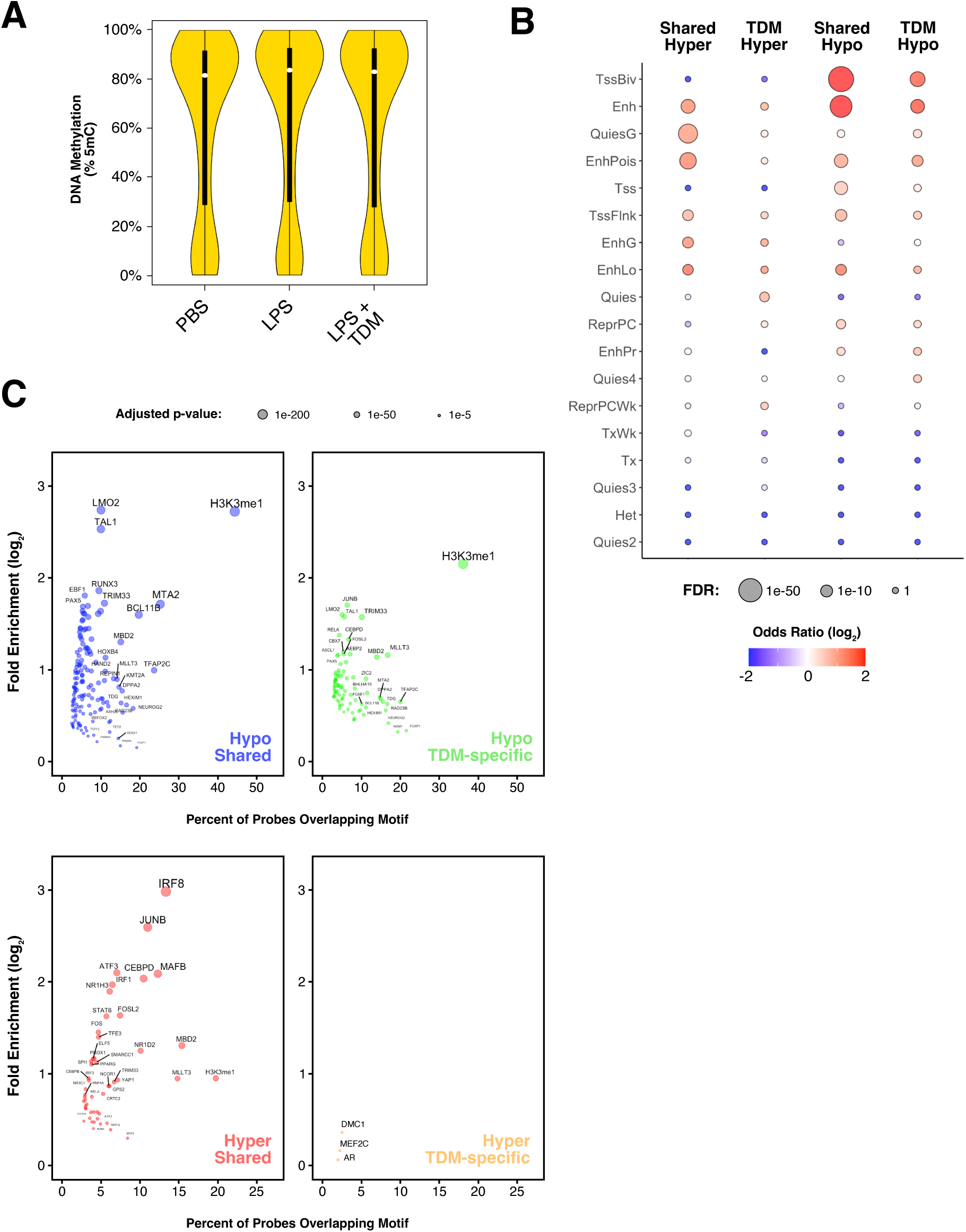
Extended analysis of DNA methylation in TDM-treated BMMCs under repetitive LPS stimulation. **(A)** Violin plots of average DNA methylation levels at each probe CpG for the BMMCs under PBS control or repetitive-LPS stimulation +/- TDM treatment, with box plots indicating medians, interquartile ranges, and 95% confidence intervals (n = 4 for each condition). **(B)** Sesame Know Your CpG (KYCG) transcription factor (TF) binding motif analysis for hyper and hypo DMRs shared between LPS and LPS+TDM cells or unique to LPS+TDM. **(C)** Enrichment of hyper and hypo DMRs in distinct chromatin states using chromHMM.

**Fig EV5.**
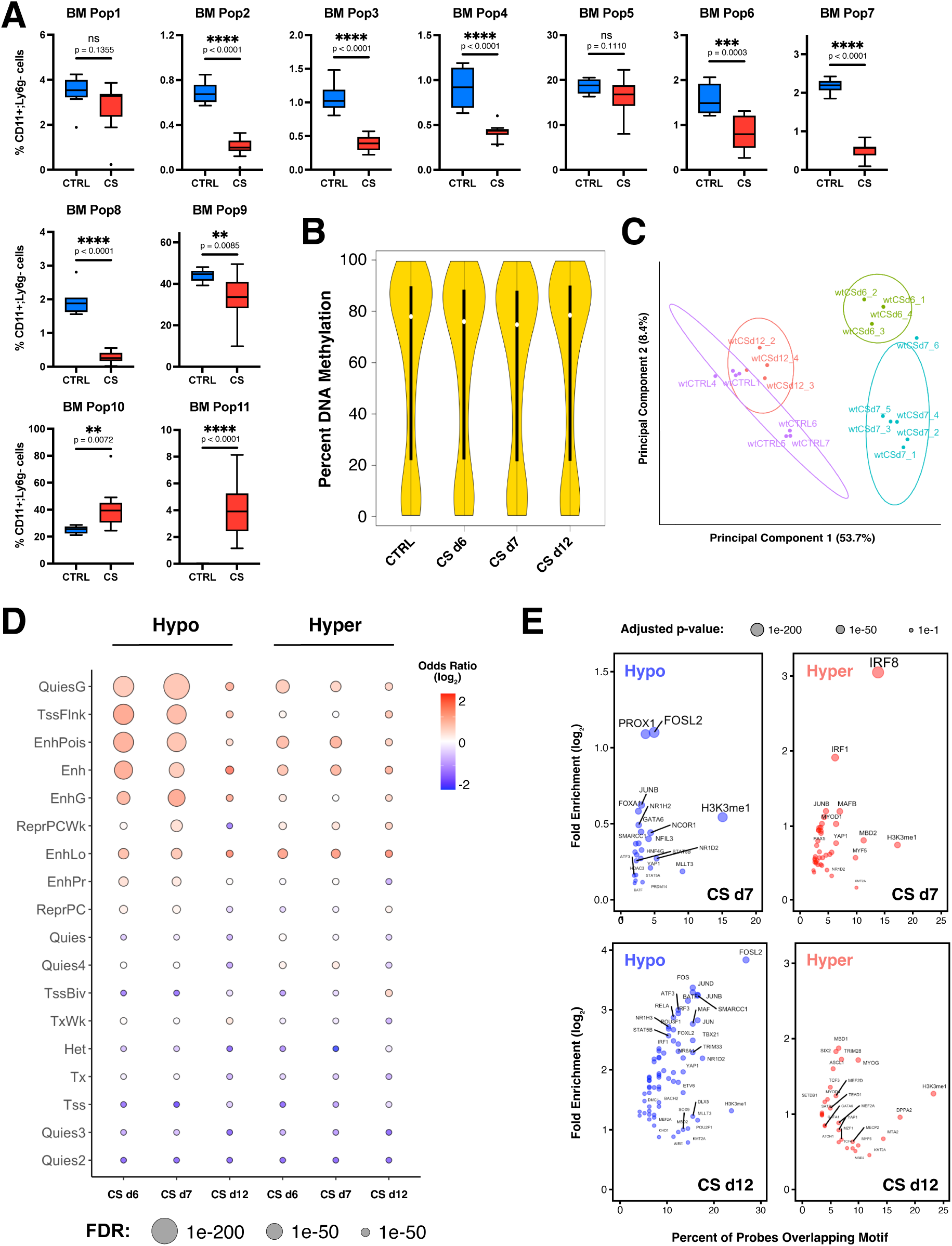
Extended analysis of *in vivo* cecal slurry flow cytometry and DNA methylation. **(A)** FlowSOM population statistics for bone marrow samples collected from control (CTRL) or cecal slurry (CS)-injected mice at experimental day 7. Box plots indicate median values (n = 8-12; two-tailed t-test; **** p-adj. < 0.0001; *** < 0.001; ** < 0.01). **(B)** Violin plots of average DNA methylation levels at each probe CpG for CTRL, CS day 6 (d6), CS d7, and CS d12 bone marrow monocyte samples, with box plots indicating medians, interquartile ranges, and 95% confidence intervals (n = 3-7 for each condition; significant hypomethylation of CS d6 and d7 relative to CTRL determined by Mood’s media test p = 3.55e-62 and 1.19e-153, respectively). **(C)** Principal component analysis of differentially methylated CpG probes in CTRL (purple), CS d6 (green), CS d7 (cyan), and CS d12 (red) bone marrow monocytes. Ovals indicate the normal distribution for each treatment. **(D)** Sesame Know Your CpG (KYCG) transcription factor (TF) binding motif analysis for hyper and hypo DMRs. **(E)** Enrichment of hyper and hypo DMRs in distinct chromatin states using chromHMM.

